# DECAPPING PROTEIN 5 regulates the circadian clock in Arabidopsis

**DOI:** 10.64898/2026.10.30.763074

**Authors:** Daniel Alejandro Careno, Soledad Perez Santangelo, Julieta Lisa Mateos, Luciana Bianchimano, Horacio Pallarés, Dominique Jacques-Vuarambon, Micaela Sol Shalom, Abril San Martín, María José de Leone, Victor Daniel Aldas Bulos, Anthony Treichel, Martin Lewinski, Rodrigo Siqueira Reis, Dorothee Staiger, Ariel Alejandro Bazzini, Marcelo Javier Yanovsky

## Abstract

The Arabidopsis transcriptome must be continuously reshaped to integrate endogenous and environmental cues, including the circadian clock and ambient light. This reprogramming relies on the coordinated synthesis and degradation of mRNAs, yet the contribution of RNA decay factors to circadian regulation remains poorly characterized. Here, we show that nearly all core clock genes are high-amplitude clock-controlled genes (HA-CCGs). A meta-analysis of transcriptome-wide mRNA decay datasets revealed that HA-CCGs have consistently shorter half-lives across multiple studies. We further identify the mRNA decapping activator DECAPPING PROTEIN 5 (DCP5) as a key regulator of circadian RNA stability. Using iCLIP, we show that DCP5 binds preferentially to HA-CCG transcripts at AU-rich motifs in their 3′ UTRs, thereby modulating their stability. Accordingly, the *dcp5* mutant displays an extended circadian period at both physiological and molecular levels. Our findings establish mRNA decay as a post-transcriptional regulatory layer that fine-tunes the decay of multiple key transcripts within the circadian system.

## Introduction

The biological clock is an endogenous mechanism that provides temporal information even in the absence of external signals and is found in organisms as diverse as bacteria, plants, and animals^1^. The clock generates biological oscillations known as circadian rhythms that regulate many aspects of an organism’s behavior, physiology, and development. The central oscillator —a group of a few dozen genes— is a gene network based on multiple interconnected feedback loops, in which a set of genes acts as mutually regulating transcriptional repressors and activators, as well as controlling the expression of thousands of other clock-controlled genes (CCG)^2^.

In Arabidopsis, the central oscillator genes *LATE ELONGATED HYPOCOTYL* (*LHY*) and *CIRCADIAN CLOCK ASSOCIATED 1* (*CCA1*) reach their peak of expression at dawn and repress the expression of the afternoon and evening genes^3–5^. Later in the morning, the *NIGHT LIGHT-INDUCIBLE AND CLOCK-REGULATED GENE* (*LNK*) family interacts with members of the *REVEILLE* (*RVE*) gene family and promotes the expression of *PSEUDO-RESPONSE REGULATOR 5* (*PRR5*), *PRR1/TIME OF CAB EXPRESSION* (*TOC1*) and *EARLY FLOWERING 4* (*ELF4*)^6–8^. Throughout the day, members of the *PRR* family, starting with *PRR9* and ending with *TOC1*, reach their respective expression peaks and sequentially suppress the expression of *LHY* and *CCA1*. In turn, *PRRs* repress one another^9,10^. After dusk, ELF3, ELF4, and LUX ARRHYTHMO (LUX) interact to form the Evening Complex (EC). The EC maintains the repression of *PRR9*, *PRR7*, *PRR5*, *GIGANTEA* (*GI*), and *LUX* itself^11–15^. The repression of *PRRs* indirectly activates the expression of *CCA1* and *LHY*, which, toward the end of the night, repress the EC, and the cycle begins again. Most of these central oscillator genes encode transcription factors and co-activators, thus, the gene circuit can explain circadian oscillations at the transcription level. However, emerging evidence suggests that mRNA circadian oscillations are also controlled in their decay phase^16^.

Two main pathways of mRNA degradation shape the stability of transcripts. These pathways begin with the conversion of a polyadenylated mRNA to an oligoadenylated mRNA through the shortening of the poly(A) tail. Then, mRNAs can be substrates of the 3’-5’ degradation pathway and be degraded by the multi-subunit complex called the exosome, or alternatively, the 5’ m^7^G cap can be removed by the decapping complex, and decapped transcripts can be degraded by the cytoplasmic 5’-3’ exoribonuclease XRN4^17^, the plant homolog of yeast and metazoan XRN1.

There is emerging evidence that shows the importance of RNA decay regulation in the function of the clock. It has been reported that alternative splicing coupled to nonsense-mediated decay controls the expression levels of *TOC1* and *ELF3*^18,19^. Also, N6-methyladenosine (m6A) modification has been shown to tune the RNA half-life of *CCA1*^20^. Regarding specific components of the RNA decay machinery, it has been shown that XRN4, and the members of the decay promoting complex LSM1-7 *SM-LIKE 1* (*LSM1*), *LSM4* and *LSM5* are required for proper clock function^21–24^.

LSM1, LSM4 and XRN4 localize in a type of membraneless cytoplasmic organelle called processing bodies (P-bodies), which provide a higher decay rate than in the rest of the cytoplasm^25^. DECAPPING PROTEIN 5 (DCP5) is a P-body component required for proper mRNA decay^26^. Here, we describe how DCP5 is necessary for the functioning of the clock, reinforcing the idea that circadian oscillations are strongly regulated at the post-transcriptional level.

## Results

### Clock-controlled transcripts have short half-lives

Circadian transcripts undergo pronounced changes in abundance over the course of a day, leading us to hypothesize that they are subject to precise temporal coordination of their transcription and mRNA degradation rates. Transcripts that oscillate with high circadian amplitude should be less stable than those with constant expression: once peak abundance is reached, the transcript must decay rapidly enough to reach its trough within ∼12 h. If certain transcripts, particularly those of the central oscillator, do not decay quickly enough, the functioning of the clock would be compromised. An extreme case can be seen when these genes are constitutively expressed using strong promoters. In these cases, a loss of rhythmicity is often observed^27–30^.

According to Romanowski *et al.* (2020)^31^, the Arabidopsis transcriptome contains 9,127 clock-controlled genes (CCGs), including both core clock genes and downstream output genes. Their relative amplitudes, defined as the amplitude normalized to the trough value, range from 0.2 to 167.3. Notably, most core oscillator genes display relative amplitudes greater than 4 (Fig. 1a). We therefore used a relative amplitude of 4 as the threshold to classify CCGs as high-amplitude (HA-CCGs, 621 genes) or low-amplitude (LA-CCGs, 8,506 genes) (Fig. 1b).

**Figure 1.**
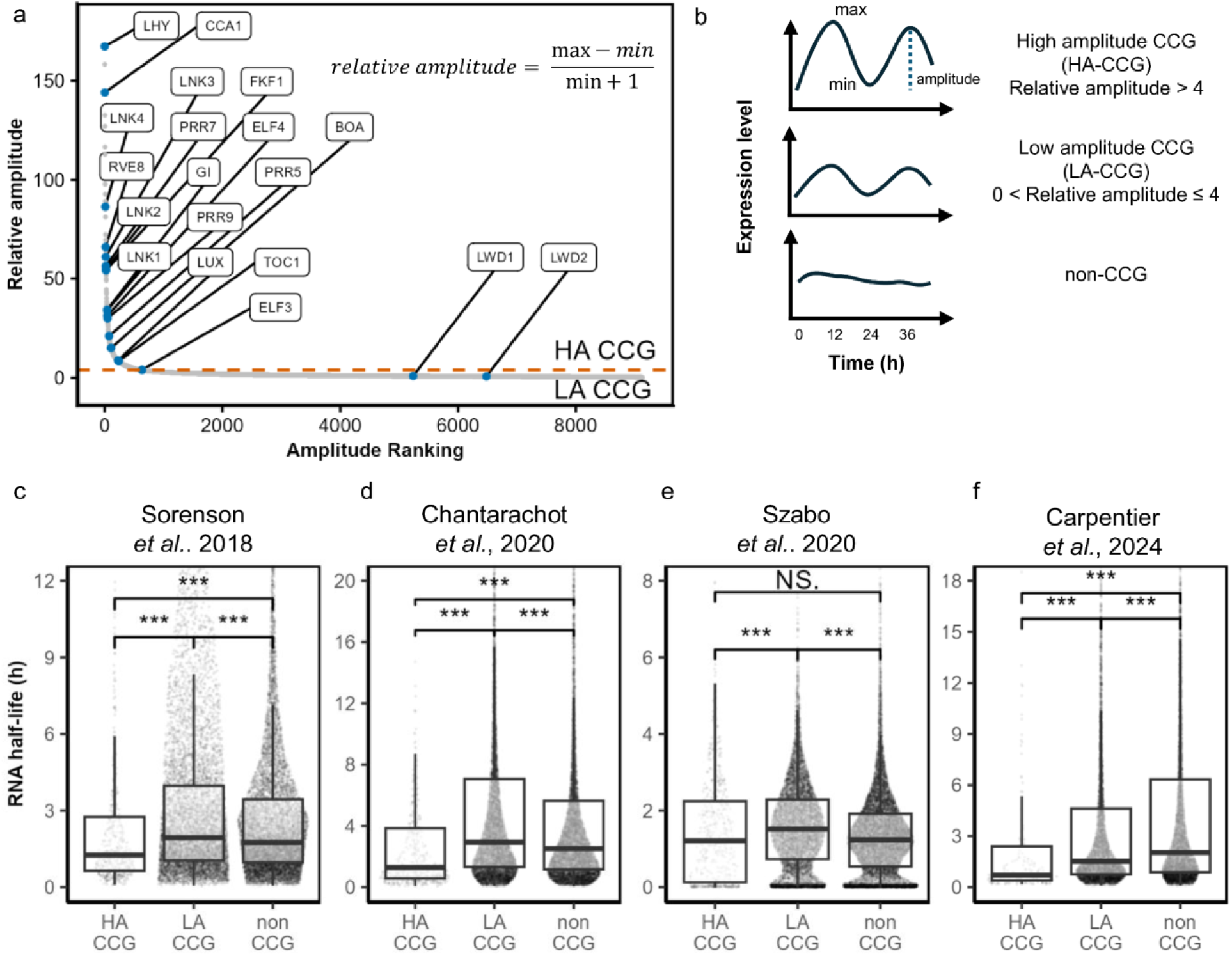
High-amplitude clock-controlled genes (CCGs) display shorter mRNA half-lives than non-oscillating genes. **a** Relative amplitude of the 9,127 CCGs identified by Romanowski *et al.* (2020)^31^ ranked in ascending order. Core oscillator genes are labeled and highlighted in green. The orange dashed line indicates the threshold at relative amplitude = 4, used to classify CCGs as high-amplitude (HA-CCGs) or low-amplitude (LA-CCGs). **b** Schematic illustrating that clock-controlled genes (CCGs) classification in HA-CCGs (relative amplitude > 4), LA-CCGs (4 ≥ relative amplitude > 0) and non-CCGs. **c-f** Distributions of mRNA half-lives for HA-CCGs, LA-CCGs, and non-CCGs across four independent genome-wide decay studies^32–35^. Statistical significance was assessed by Kruskal–Wallis test followed by pairwise Wilcoxon rank-sum tests with Benjamini–Hochberg (FDR) correction; *P < 0.05, **P < 0.01, ***P < 0.001; NS, not significant.

To test our hypothesis, the half-lives of CCGs were compared based on their amplitude using previously published transcriptome-wide datasets reporting half-lives for Arabidopsis mRNAs^32–35^. In all four datasets, the median half-life of HA-CCG is lower than that of both LA-CCG and non-CCG, and in three cases out of the four these differences are statistically significant (Fig. 1c-f). Furthermore, when CCGs were grouped into amplitude deciles, half-lives trended downward with increasing amplitude (Supplementary Fig. 1). These observations align with microarray data showing enrichment of CCGs among fast-decay transcripts^36^. The consistently shorter half-lives of HA-CCG mRNAs across independent datasets suggest that post-transcriptional control of RNA stability plays a key role in shaping circadian oscillations.

### DCP5 is required for normal clock function

Previously, two components of the 5’-3’ RNA decay pathway, LSM1 and XRN4, have been described as necessary for correct clock function^22,23^. These RNA decay factors may be part of a mechanism that maintains CCG mRNA half-lives within a range compatible with proper circadian oscillations, that is, the appropriate period, phase, and amplitude. The 5’-3’ mRNA decay pathway requires removal of the 5’ m^7^G cap by the decapping complex. In Arabidopsis, the core decapping complex includes DCP1, DCP2 (the component with enzymatic activity) and VARICOSE (VCS), and its assembly is modulated by DECAPPING PROTEIN 5 (DCP5)^37,38^. While *dcp1*, *dcp2* and *vcs* are all post-embryonic lethal mutants^39^, a hypomorphic *dcp5* mutant allele, *dcp5-1*, can grow and complete its developmental life cycle^26^. We therefore studied clock function in the *dcp5-1* mutant and found a long circadian period phenotype for the rhythm of leaf movement in *dcp5* compared with wild-type plants (24.83 ± 0.44 h vs 27.57 ± 0.72 h) (Fig. 2a-b). Supporting the idea that the circadian phenotype observed in this mutant is due to the mutation in *DCP5*, this phenotype was completely rescued when we complemented *dcp5-1* plants with a WT version of the *DCP5* gene (Fig. 2b). We then analyzed circadian rhythms in gene expression by evaluating rhythms in bioluminescence driven by the *pTOC1:Luc* and *pCCR2:Luc* reporter genes, and we also observed a strong circadian phenotype in *dcp5-1* mutant plants. Indeed, the rhythm of expression of both reporters was lengthened by ∼2 h in *dcp5-1* mutants compared with WT plants (24.18 ± 0.57 h vs 26.28 ± 0.34 h for *pTOC1:Luc*, and 23.65± 0.48 h vs 25.59 ± 0.50 h for *pCCR2:Luc*) (Fig. 2c-d and Supplementary Fig. 2). Together, these results demonstrate that DCP5 is required for proper clock function and reinforce the view that mRNA decay constitutes a key regulatory layer for circadian rhythms.

**Figure 2.**
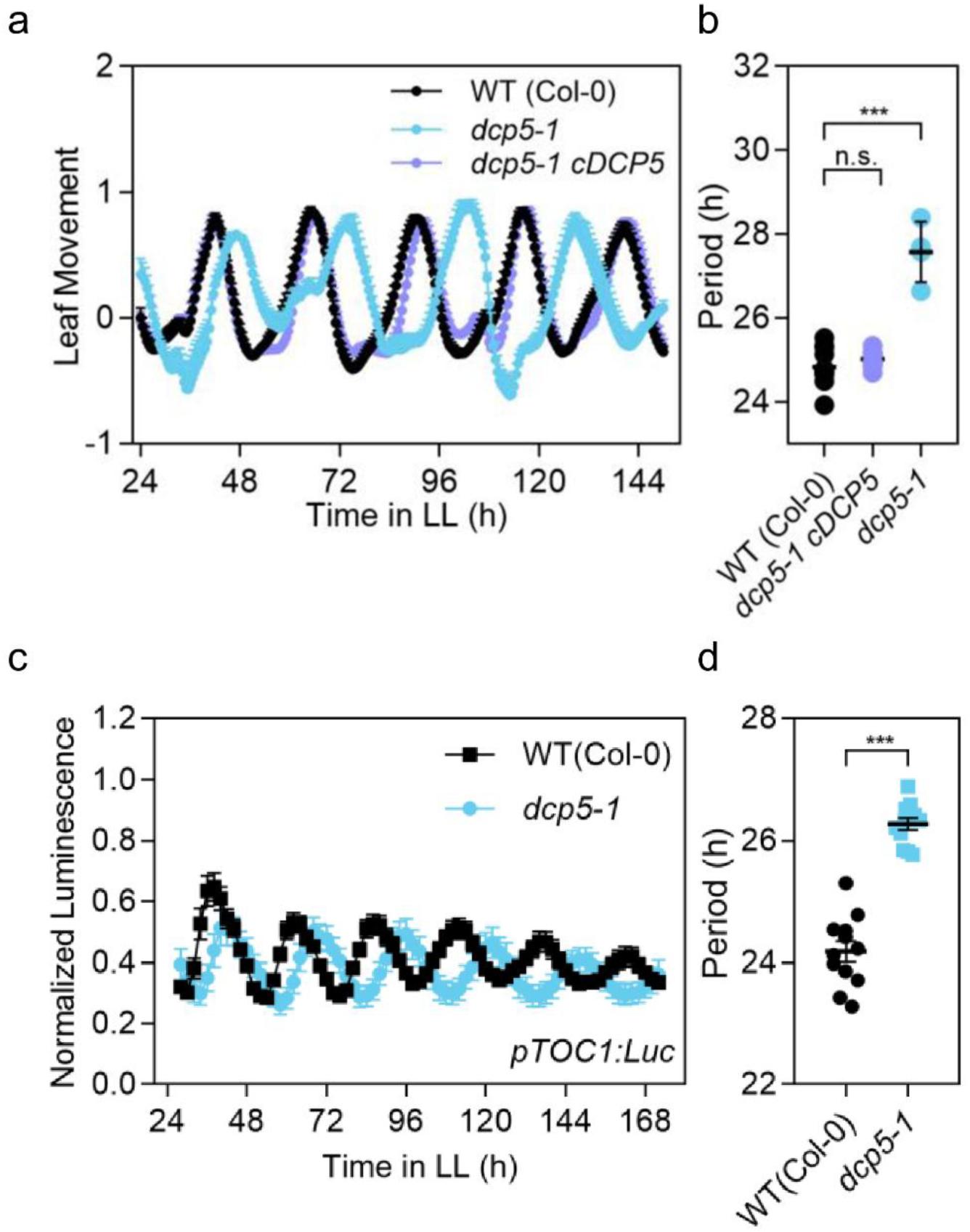
DCP5 is required for proper clock function. **a-b** Circadian rhythm of leaf movement. **(a)** Leaf angle and **(b)** period length estimation of entrained under LD conditions and then transferred to LL. WT (Col-0) (n=12), *dcp5-1* mutant (n=4), *dcp5-1 DCP5* (n=4). **c-d** Luciferase reporter assay. **(c)** Expression of the night core-clock reporter *pTOC1:Luc* and **(d)** period length estimation. The expression was measured for 6 d in LL after entrainment under LD conditions. WT (Col-0) (n=12), *dcp5-1* mutant (n=12). The length of leaf movement rhythms and bioluminescence was estimated by the fast Fourier transform nonlinear least squares (FFT-NLLS) test. Data are mean ± s.e.m. ***P < 0.001 (One-way ANOVA and two-tailed Student’s t-test).

### DCP5 is a global regulator of RNA stability

To understand how DCP5 shapes the RNA decay landscape, we performed a cordycepin transcriptional arrest assay followed by whole-transcriptome RNA sequencing. Wild-type and *dcp5* plants were treated with cordycepin, and samples were collected 0, 30, 60, and 90 min later. After sequencing, transcript abundances were used to estimate mRNA decay rates in each genotype. RNA half-lives spanned a wide range, as previously reported^32^, with median values of 60 min and 67 min in wild-type and *dcp5* plants, respectively. Among the genes with the fastest decay (half-life < 10 min), three belonged to the *SMALL AUXIN UP REGULATED* (*SAUR*) gene family (AT2G21210, AT4G38840, and AT4G31320), which have previously been described as short-lived^40^. Of the 13,698 genes that passed the minimum expression threshold and met the model-fitting criteria, 3,206 were more stable in *dcp5* plants, whereas 1,435 were more stable in wild-type plants. Loss of DCP5 mainly affects genes with the shortest half-lives, with 80% of those genes having a half-life shorter than 90 min (Fig. 3a). A Gene Ontology (GO) term enrichment analysis revealed that categories related to mRNA translation, photosynthesis and response to environmental stimuli, which are in part regulated by the biological clock, were overrepresented among transcripts stabilized in the *dcp5* mutant (Fig. 3b).

**Figure 3.**
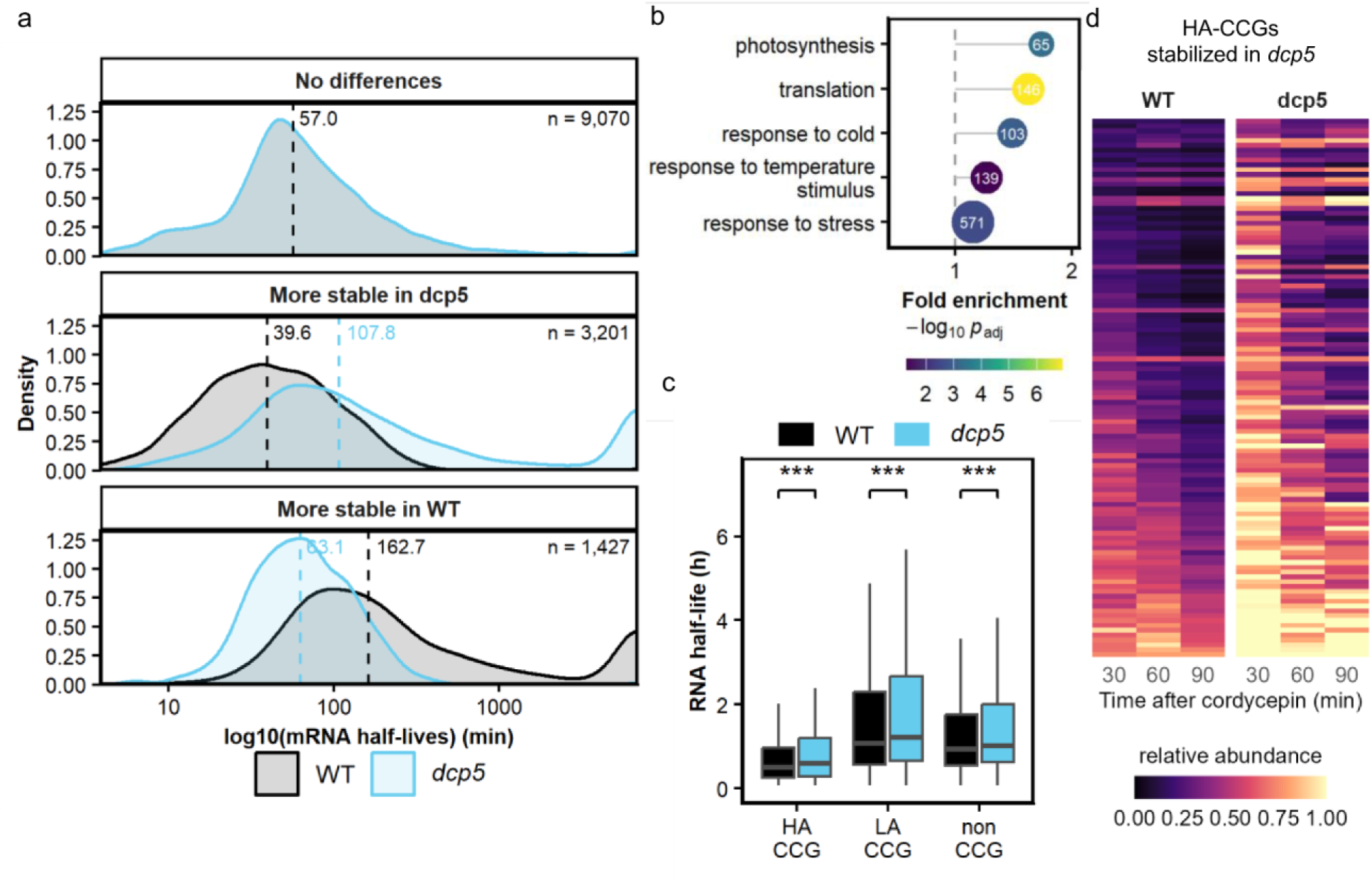
DCP5 controls mRNA half-lives globally. **a** Distribution of mRNA half-lives in WT and *dcp5* for transcripts stabilized in *dcp5* (n = 3,206), stabilized in WT (n = 1,435) and with unchanged stability (n = 9,070). The dashed lines mark the mean half-life in each genotype. Genes with half-lives of 6,931.47 min (α = 0.0001 min⁻¹, the lower bound of the model) in both genotypes were excluded. **b** Gene Ontology (GO) term (biological process) enrichment analysis of transcripts stabilized in the *dcp5* mutant. Manually selected terms are shown. X-axis position gives fold enrichment (observed/expected number of genes); the dashed line marks no enrichment (fold = 1). Bubble color indicates significance (−log₁₀ adjusted P; one-sided hypergeometric test, Benjamini–Hochberg correction); the number within each point is the count of genes from the list annotated to that term. **c** RNA half-lives distribution across high amplitude (HA-), low amplitude (LA-), and non-clock-controlled genes (CCGs) in WT and *dcp5* plants. Within each group, WT and *dcp5* half-lives were compared gene by gene using a two-sided Wilcoxon signed-rank test, with P values adjusted across the three groups by the Benjamini–Hochberg method (*** adjusted P < 0.001) **d** Decay profiles of HA-CCGs stabilized in *dcp5* (n = 111). Heatmaps show transcript abundance at 30, 60 and 90 min after cordycepin treatment, relative to t0, for WT (left) and *dcp5* (right). Rows are genes, in the same order in both panels, sorted by WT half-life (shortest at the top).

We found that the half-life tendencies of CCG mRNAs previously observed in other datasets (Fig. 1c-f and Supplementary Fig. 1) were conserved in our dataset (Fig. 3c; Supplementary Fig. 3). Consistent with the clock phenotype observed in the *dcp5* mutant (Fig. 2 and Supplementary Fig. 2), CCGs decayed more slowly in the mutant (Fig. 3c). Among HA-CCGs, 280 showed no difference between genotypes and 26 were stabilized in WT (Supplementary Fig. 4), while 111 had longer half-lives in *dcp5* (Fig. 3d). Notably, only a few core clock genes showed differences in their decay. This was also observed in the *xrn4* mutant^22^, suggesting that the clock phenotypes observed in RNA decay mutants may reflect coordinated changes in the decay of multiple clock-associated transcripts rather than effects restricted to a small subset of core clock genes.

### High-amplitude transcripts preferentially bind to DCP5

To identify the RNAs that are bound by DCP5 *in vivo* and the specific binding nucleotide sequence we performed individual-nucleotide resolution UV crosslinking and immunoprecipitation (iCLIP). We collected samples of *dcp5* seedlings expressing DCP5-YFP-HA at two time points to capture CCG transcripts across the circadian cycle (Fig. 4a). We detected 96,632 binding sites in 9,116 genes, with 67.7% of targets detected at both time points (Fig. 4b). Although the number of transcripts that were present in both datasets was high, some degree of time specificity was evident when we examined those transcripts with expression peaks at CT12 and CT24 (Supplementary Figs. 5 and 6). A similar high overlap but also time-of-day dependent targets was observed for the clock-regulated RNA-binding protein *At*GRP7 at CT24, the circadian minimum, and at CT36, the circadian maximum of its expression^41^.

**Figure 4.**
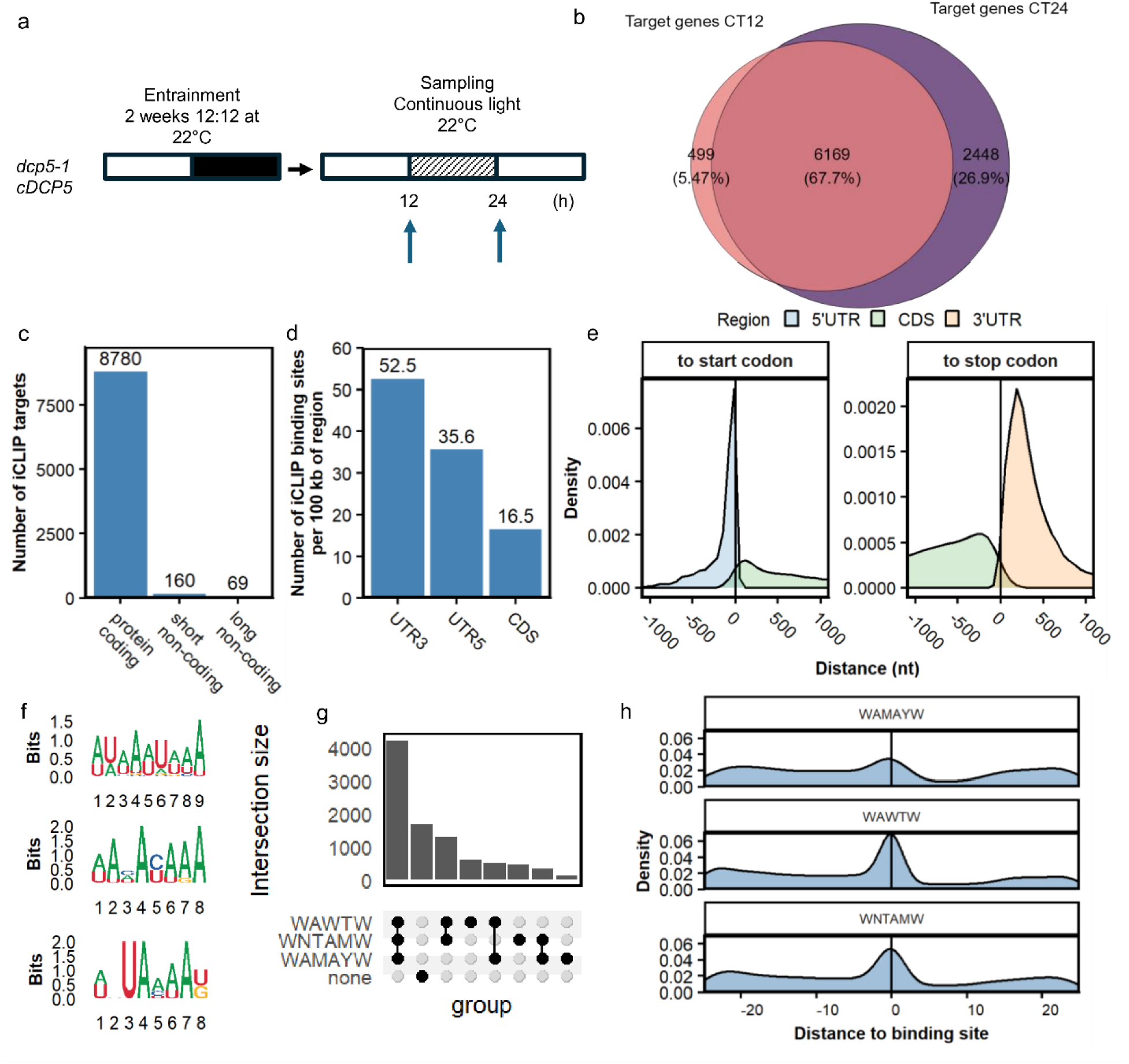
DCP5 binds preferentially protein-coding RNAs in their 3’UTR. **a** Experimental design. *dcp5-1* plants expressing DCP5-YFP-HA were entrained under 12 h light/12 h dark cycles at 22 °C and transferred to continuous light. Samples were collected at subjective dusk (CT12) and subjective dawn (CT24) for iCLIP. **b** Overlap between genes with DCP5 binding sites at CT12 and CT24. **c** Number of DCP5 target genes by biotype (CT12 and CT24 combined). Short non-coding includes snoRNA, tRNA, snRNA, miRNA and rRNA. Long non-coding includes lncRNA and other ncRNA. **d** Density of DCP5 binding sites in 5′ UTRs, CDSs and 3′ UTRs, as binding sites per 100 kb of the total length of each region. Both time points combined. **e** Distribution of distances (nt) from binding sites to the start codon (5′ UTR and CDS sites) or to the stop codon (CDS and 3′ UTR sites). Negative values are upstream of the respective codon. **f** Sequence logos of the three top-scoring motifs found by SSMART in 64 nt windows centered on binding sites (both time points combined). **g** UpSet plot of DCP5 target genes grouped by which motifs occur in their binding sites. None: genes with no binding site matching any of the three motifs. Intersections with fewer than 10 genes are not shown. **h** Distribution of distances (nt) between each motif occurrence shown in (f) and the nearest DCP5 binding site within 25 nt. Positive values mean the motif is downstream of the binding site. (IUPAC code, W = A/U; M = A/C; Y = C/U).

Consistent with its role in mRNA decay, 97% of DCP5 targets were protein-coding, a significant overrepresentation relative to the 84% of protein-coding genes in the Arabidopsis genome (p < 0.001, hypergeometric test) (Fig. 4c). Binding sites were preferentially located in the 3’UTR and only 8.5% (772 genes) of all target transcripts lacked a binding site in the 3’UTR (Fig. 4d, Supplementary Fig. 7). 5′UTR sites peaked ∼2 nt upstream of the start codon, whereas 3′UTR sites peaked ∼220 nt downstream of the stop codon, typically near the 3′ end of the transcript (Fig. 4e, Supplementary Fig. 8). A motif search within a 28 nt window on either side of each binding site revealed enrichment for AU-rich sequences (Fig. 4f). Of all DCP5 targets, 85.8% contained at least one of these motifs near the binding site (Fig. 4g). The binding sites usually overlapped with these motifs (Fig. 4h). A transcriptome-wide search, across both DCP5-bound and unbound transcripts, showed that these motifs were most densely distributed within coding sequences (CDS) (Supplementary Fig. 9). However, DCP5 bound these motifs preferentially when they were located in the 3′UTR (Supplementary Fig. 9), suggesting that additional trans-acting factors are required to recruit DCP5.

The half-lives of DCP5 targets had a median value of 59 min in WT and 72 min in *dcp5* (paired Wilcoxon signed-rank test, P < 0.001), while the difference in the half-lives of non-targets was less pronounced, with a median of 61 min and 64 min in WT and *dcp5* (paired Wilcoxon signed-rank test, P < 0.001), respectively (Supplementary Fig. 10). Notably, DCP5 bound the transcripts of most HA-CCGs (chi-square test, p < 0.001) (Fig. 5a). These targets included most core clock genes. Moreover, both the binding site density and the PureCLIP scores, which reflect the confidence and strength of individual binding sites, were higher in HA-CCGs than in LA-CCGs and non-CCGs (Fig. 5b, c). These results suggest that DCP5 binds and likely regulates the stability of multiple clock gene transcripts and many of their downstream circadian output genes.

**Figure 5.**
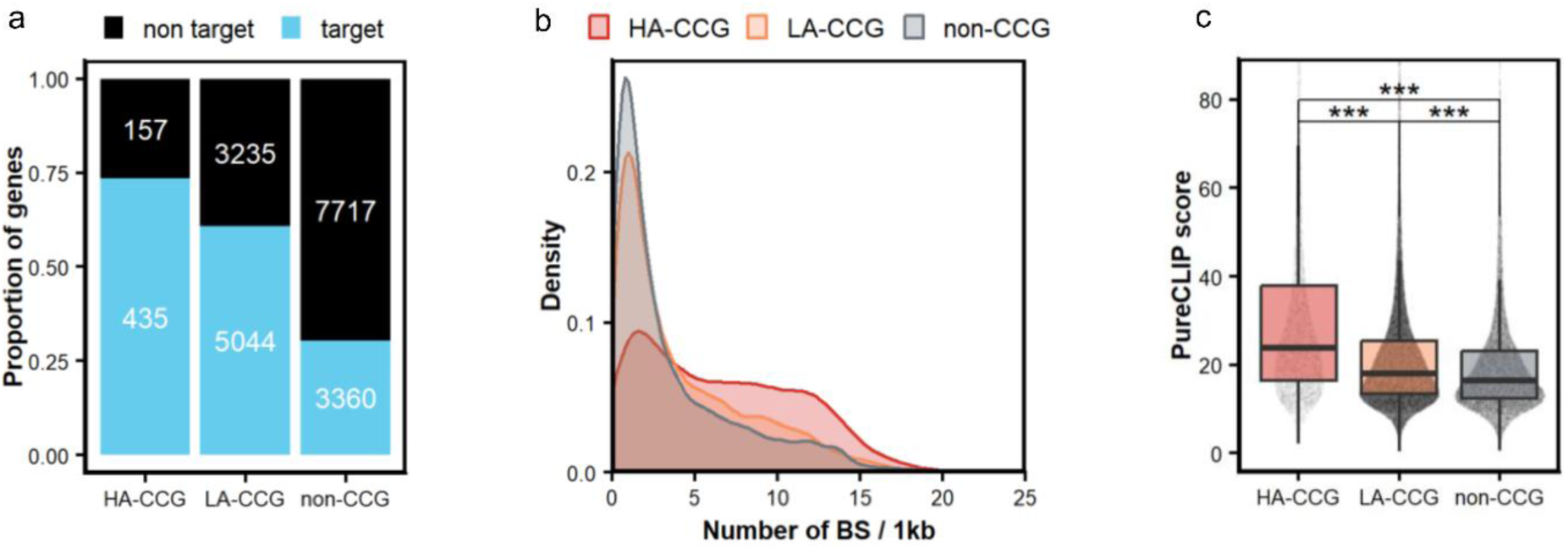
High-amplitude clock-controlled genes (HA-CCG) are overrepresented among DCP5 targets. **a** Proportion of HA-CCG, LA-CCG and non-CCGs that are targeted by DCP5 (chi-square test, p < 0.001). CCGs and their relative amplitudes are from Romanowski *et al*. (2020). **b** Distribution of binding-site density (binding sites per kb of the longest transcript) in DCP5 target genes, by CCG category. **c** PureCLIP scores of DCP5 binding sites in HA-CCGs (n = 6,936 sites in 447 genes), LA-CCGs (n = 60,119 sites in 5,047 genes) and non-CCGs (n = 29,577 sites in 3,622 genes), CT12 and CT24 combined. Kruskal–Wallis test followed by Dunn’s test with Benjamini–Hochberg correction; ***, adjusted P < 0.001.

### DCP5 regulates at least 25% of the transcriptome

To assess the impact of DCP5 on the transcriptome, we performed RNA-seq on wild-type and *dcp5* plants collected every 4 h over one day in free-running conditions after entrainment under a 12 h light/12 h dark photoperiod (Fig. 6a). We then applied the Jonckheere-Terpstra-Kendall (JTK) algorithm to identify genes undergoing circadian oscillations, detecting 8,036 CCGs in WT (JTK BH.Q < 0.01). Of these, 5,399 genes (67%) overlapped with CCGs identified by Romanowski *et al.* (2020)^31^, with a relative amplitude Pearson correlation of 0.83 (Fig. 6b, Supplementary Fig. 11a). Also, we observed a distribution of relative amplitudes comparable to that previously observed in the dataset from Romanowski *et al.* (2020)^31^, with most core clock genes classified as HA-CCGs (Supplementary Fig. 11b).

**Figure 6.**
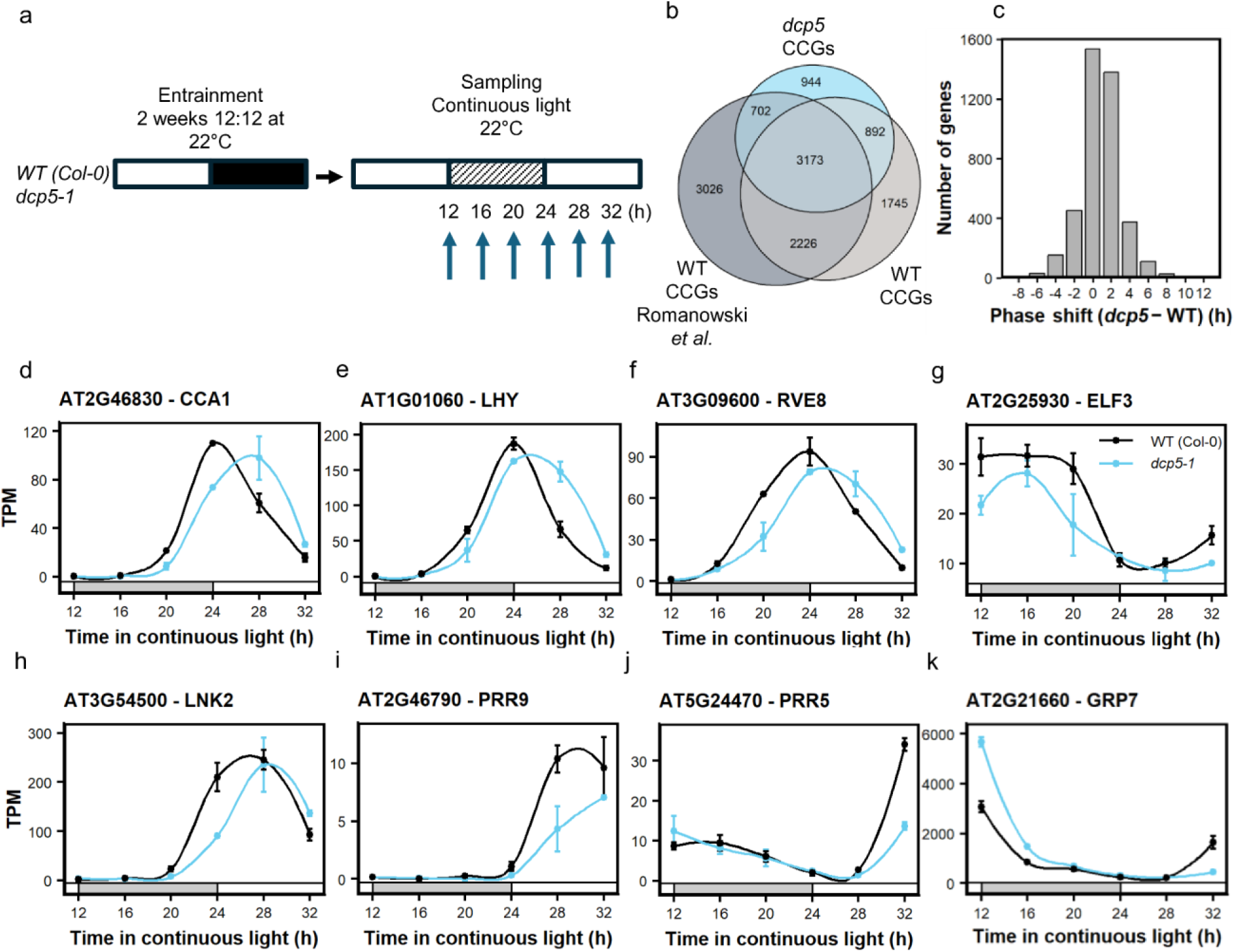
The circadian transcriptome needs DCP5 for a correct time of peak expression. **a** Experimental design. WT (Col-0) and *dcp5-1* seedlings were entrained under 12 h light:12 h dark cycles at 22 °C and then moved to continuous light. Samples were collected every 4 h from CT12 to CT32 (six time points; n = 2 biological replicates). **b** Euler diagram showing how circadian clock-regulated genes (CCGs) overlap between three sets: WT and *dcp5-1* in this study, and WT in Romanowski *et al.* (2020). In this study, CCGs were identified with JTK_CYCLE (BH-adjusted q < 0.01). **c** Peak phase of each gene in WT versus *dcp5-1*, estimated with JTK_CYCLE. Only genes rhythmic in both genotypes are shown (BH-adjusted q < 0.01; n = 4,065). **d-k** Expression of representative clock genes **d** *CCA1*, **e** *LHY*, **f** *RVE8*, **g** *ELF3*, **h** *LNK2*, **i** *PRR9*, **j** *PRR5* and **k** *GRP7*. Points show mean TPM ± s.e.m. (n = 2), and lines are loess fits. Gray and white bars mark subjective night and subjective day.

We also detected 5,711 CCGs in *dcp5*, including 4,065 genes (71%) that were also detected as CCGs in our WT dataset (Fig. 6b). The Pearson correlation between relative amplitudes in WT and *dcp5* was 0.89, suggesting that DCP5 does not substantially alter the relative amplitude pattern across genes (Supplementary Figure 12a). However, we observed similar phase shifts in 62% of shared CCGs (Fig. 6c, Supplementary Fig. 12b). These shifts are evident in the expression patterns of some core clock genes in *dcp5* (Fig. 6d-g), which are consistent with the lengthened period observed in the leaf movement and luciferase reporter assays (Fig. 2).

To further understand the effects of DCP5 on the transcriptome, we performed a differential expression analysis. We defined differentially expressed genes (DEGs) between WT and *dcp5* as genes whose expression was significantly affected by genotype across the time course, either at all time points or in a time-of-day-dependent manner (likelihood-ratio test, adjusted P < 0.05). DEGs that showed a change of at least 1.5-fold at one or more time points (adjusted P < 0.05) were classified as “robust”, and the remainder as “soft” (See Methods). Thus, we identified 801 robust and 5,964 soft DEGs. The subset of robust DEGs showed the highest proportions of DCP5 targets, transcript stabilized in *dcp5*, and HA-CCGs, while the subset of soft DEGs showed larger proportions in those categories than non-DEGs (Supplementary Fig. 13). Among robust DEGs, only 27% (220 out of 801) showed an expression difference of |log2FoldChange| > log2(1.5) at two or more time points. This suggests the DCP5 effect is time-dependent. Notably, most core clock genes (17 of 27) were differentially expressed (4 soft and 13 robust DEGs).

Altogether, these findings suggest that loss of DCP5 does not produce large changes in overall transcript levels but instead alters temporal expression patterns throughout the day. We identified a subset of genes with altered circadian expression patterns that was enriched for DCP5 targets. These HA-CCGs do not reach higher peak expression levels in the *dcp5* mutant but instead exhibit a shift in the time of peak expression, consistent with a longer period, suggesting that DCP5 contributes to the decline in transcript abundance after peak expression.

## Discussion

Here we show that the mRNA decapping activator DCP5 is a key component to maintain a correct expression phase of transcripts in the circadian network in Arabidopsis. Using iCLIP, we found that DCP5 binds roughly one-third of the transcriptome, with marked enrichment for high-amplitude clock-controlled genes (CCGs). The *dcp5* mutant displayed long-period rhythms in leaf movement and clock-gene expression, together with a global delay in the phase of peak expression of CCGs. DCP5 targets were more stable in *dcp5*, consistent with a role for DCP5 in promoting mRNA decay. The number of transcripts stabilized in *dcp5* is smaller than that in other decay mutants such as *vcs* or *rh6/8/12*^32,33^. This could be due to differences in experimental design and may also be related to the lethal phenotypes associated with the loss of VCS or the combined loss of RH6, RH8, and RH12^33,39,42^. A small subset of targets instead had shorter half-lives in the mutant, which could reflect an indirect effect whereby accumulated transcripts are channeled into the RNA quality control small interfering RNA (rqc-siRNA) pathway^43,44^.

The long-period phenotype of *dcp5* mirrors the clock defects of other RNA-turnover mutants, including *xrn4* and *lsm1*^22,23^. The generally milder phenotypes of *xrn4* compared with *dcp5* and *lsm1* suggest a hierarchical organization in which DCP5 and LSM1 act upstream of XRN4. By sequestering and translationally repressing target mRNAs, DCP5 and LSM1 may commit transcripts to translational repression even in the absence of XRN4. In *dcp5* and *lsm1* mutants, by contrast, XRN4 would both lose this trigger and degrade its substrates less efficiently, because they are no longer concentrated in cytoplasmic granules. Similarly, the *dst1* mutant, in which clock transcripts have longer half-lives, also has a long period^45^. In contrast, loss of the cap-binding factor eIF4E, which shields transcripts from degradation^46^, shortens the circadian period^47^. Likewise, in Drosophila, knockdown of the translation-promoting factor LARK shortens the period^48^. Together, these observations support a model in which impaired decay and mRNA accumulation lengthen the period (Fig. 7), whereas loss of mRNA stabilization accelerates the clock.

**Figure 7.**
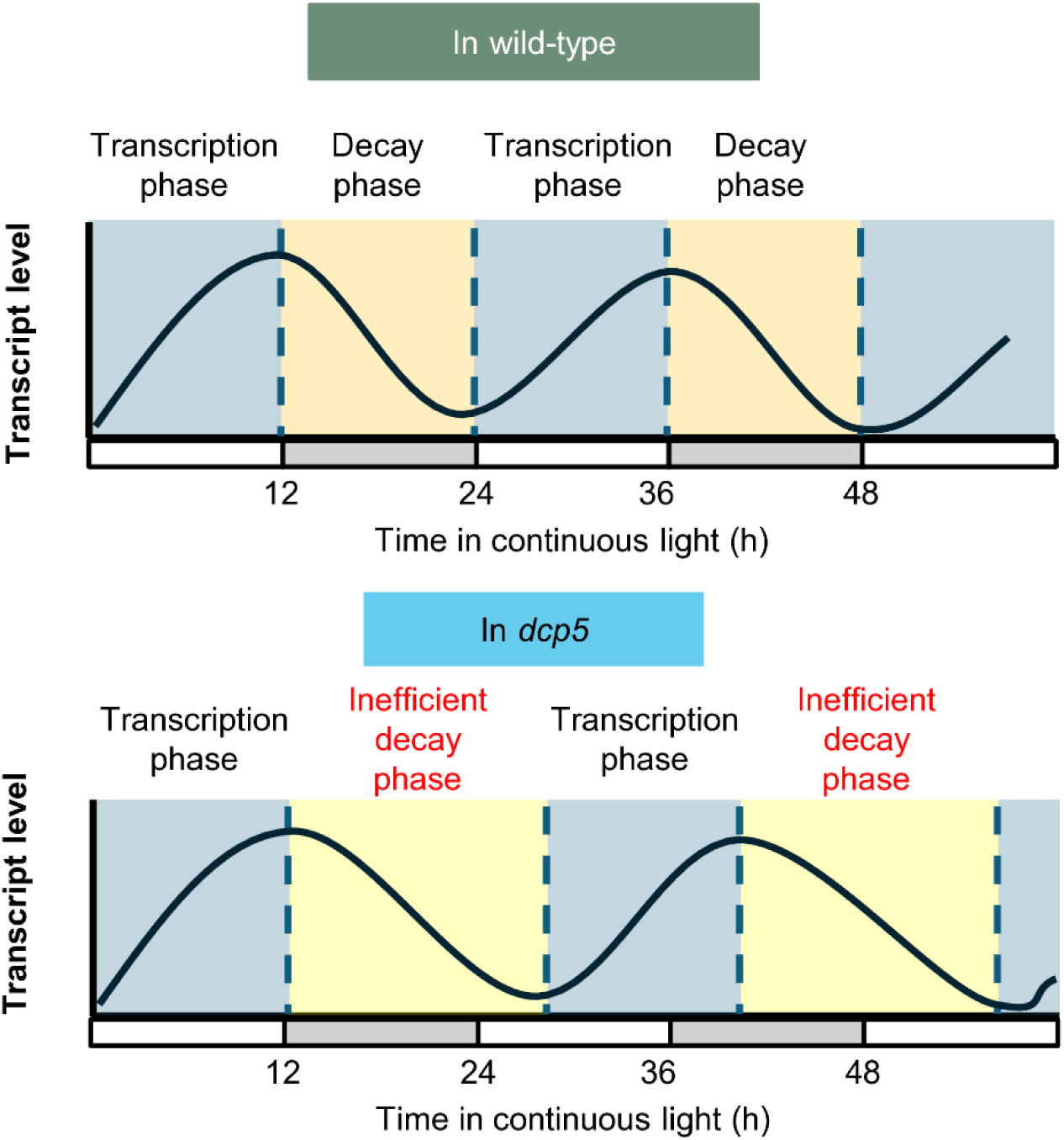
Proposed role of RNA decay machinery in pacing circadian oscillations. In WT after the peak of expression is reached, mRNA enters a decreasing phase until it reaches its minimum. When the decay machinery is impaired, such as in the *dcp5* mutant, the mRNA degradation is slower resulting in an extended circadian period.

Interestingly, DCP5 transcript levels do not oscillate, suggesting that *DCP5* expression is not under clock control. However, DCP5 interacts with many RNA-binding proteins whose mRNAs do oscillate^31,49^, raising the possibility that its activity or target specificity is modulated rhythmically through its interactome. Also, it has been reported that DCP5 physically interacts with multiple poly(A) binding proteins (PAB2, PAB3, PAB4, PAB5 and PAB8)^49^, which is consistent with DCP5 preference for binding at the end of 3’ UTRs. Besides its role in the circadian clock, DCP5 is involved in multiple physiological processes such as photomorphogenesis^50^, response to osmotic stress^51,52^ and response to hormones^53^. All these processes presumably require a re-programming of the transcriptome, in which the DCP5 interactome could have a major role in selecting its target transcripts.

More broadly, our results identify DCP5 as a post-transcriptional hub through which control of mRNA stability shapes clock timing. A constitutively expressed decapping activator whose activity is tuned by a rhythmic interactome could provide robust yet flexible control over large sets of transcripts. An important open question is the extent to which this regulation propagates to the translatome and proteome, particularly given that a substantial fraction of rhythmic proteins arise from non-rhythmic transcripts^54^. Addressing this question will be central to understanding how RNA turnover contributes to rhythmic physiology in plants.

## Methods

### mRNA half-life meta-analysis

Half-lives were extracted from published Supplementary Tables. When needed, half-lives were estimated from decay rate constant (alpha) as t_1/2_=ln(2)/alpha. In the case of Szabo *et al.* (2020) compound model estimations were used.

### Plants and growth conditions

Plants were grown in pots containing a mixture of organic substrate, vermiculite, and perlite [0.5:1:1 (vol/vol)] at the particular photoperiod and temperature conditions indicated in each experiment. The mutant *dcp5-1* (SALK_106536) was obtained from the Arabidopsis Biological Resource Center (ABRC).

### Generation of dcp5 DCP5 transgenic plants

DCP5 CDS was cloned from WT cDNA and introduced in pDONR/Zeo vector (Invitrogen) via the Gateway method. A binary vector containing this fragment was generated by LR recombination with pEarlyGate 101, thus obtaining the DCP5-YFP-HA protein under the control of the CaMV35S promoter. This plasmid was introduced into *Agrobacterium tumefaciens* GV3101 strain and transformed into *dcp5-1* plants using the floral dip method ^55^. Transgenic lines were selected on Murashige and Skoog media supplemented with 15 mg/L PPT (Duchefa).

### Leaf movement assay

For leaf movement analysis, plants were entrained under a long-day photoperiod (16 h light/8 h dark cycles, 80 μmol m⁻² s⁻¹), transferred to continuous white light, and the angle of the first leaf pair was recorded every 1 h for 5–6 days using ImageJ (imagej.nih.gov/ij/). Period estimates were calculated using BioDare2 (biodare2.ed.ac.uk) and analyzed by FFT-NLLS. Statistical analysis was performed using a two-tailed Student’s t-test or a one-way ANOVA, as appropriate.

### Luciferase reporter assays

Seedlings were grown on Murashige and Skoog medium with 0.8% agar supplemented with 1% sucrose in a 96-well plate. One seed was placed per well, and the plate was kept in darkness at 4 °C for 3 days. The seedlings were then entrained under a long-day cycle (16 h light/8 h dark, 80 μmol m⁻² s⁻¹). After 7 days, luciferin was added to each plant, the plate was transferred to constant light (50 μmol m⁻² s⁻¹) and temperature conditions (22 °C) and placed in a luminometer to measure the bioluminescence emitted by each seedling every 2 h. After 5–6 days, data analysis was performed using Mikrowin 2000 software (version 4.29) (Mikrotek Laborsysteme). The luminometer used was the Centro LB-960 microplate reader (Berthold Technologies). Period estimates were calculated using BioDare2 (biodare2.ed.ac.uk) and analyzed by FFT-NLLS. Statistical analysis was performed using a two-tailed Student’s t-test.

### Hypocotyl elongation assays

Seeds were sown in pots, stratified for 2 days at 4 °C in darkness, and then grown in a 12 h light/ 12 h of dark photoperiod at 22 °C (80 μmol m⁻² s⁻¹). After germination, plants were kept either in the same condition or transferred to the same photoperiod (80 μmol m⁻² s⁻¹) with 28 °C days and 22 °C nights. Photographs of the plants were taken every 1 h and the hypocotyls were measured using ImageJ (imagej.nih.gov/ij/).

### Cordycepin time course experiment and mRNA decay analysis

Plants were grown in continuous light (LL, 50 μmol m⁻² s⁻¹) from germination, with no entrainment. After 2 weeks, seedlings were transferred to flasks containing a buffer (1mM 1,4-piperazinediethanesulfonic acid, pH 6.25, 1mM sodium citrate, 1mM KCl and 15mM sucrose), and after a 30-min incubation, cordycepin (0.6mM) was added, vacuum infiltration was performed and the samples were taken at 0, 30, 60 and 90min. Total RNA was extracted using TRIzol reagent (Invitrogen) and samples were stored for shipment in GenTegra RNA tubes. Sequencing libraries were generated using Abclonal Fast RNA-seq Lib Prep Kit V2 (RK20306) with polyA capture by VAHTS mRNA Capture Beads (Cat.No.N401-01/02) and sequenced on the Illumina NovaSeq X Plus platform, providing 150-bp paired-end reads (Novogene, Singapore).

The RNA sequencing reads were mapped to the Arabidopsis thaliana TAIR10 genome^56^ using STAR (version 2.7.11)^57^ with the following parameters: --twopassMode Basic --outFilterMultimapNmax 2 --outFilterType BySJout --outSJfilterReads Unique --sjdbOverhang 149 --alignSJoverhangMin 6 --alignSJDBoverhangMin 3 --alignIntronMin 20 --alignIntronMax 5000.

Modeling of RNA decay was performed as previously reported^32,33^. Briefly, each transcript’s TPM expression was normalized to its genotype-specific mean T0 value and then corrected by a decay factor estimated separately for each genotype × time point condition. Decay factors were derived from reference genes selected as those that (i) were among the 500 most highly expressed genes at T0, (ii) showed no significant expression difference between genotypes at T0 (excluded if padj < 0.05 and |log2FC| ≥ 1), and (iii) showed monotonically increasing T0-normalized expression across all four time points (T0 < T30 < T60 < T90) in both WT and*dcp5*. Using those criteria, we found and used for the decay factors 37 reference genes: AT3G09260, AT5G26000, AT1G54410, AT5G35630, AT1G23130, AT4G04640, AT4G37930, AT5G42980, AT5G25980, AT3G14420, AT1G23310, AT1G02780, AT1G62380, AT1G11860, AT2G36530, AT1G20020, AT2G30950, AT1G78630, AT3G14415, AT1G43170, AT5G44340, AT5G09660, AT4G02520, AT1G09690, AT1G09340, AT5G39570, AT3G11940, AT5G30510, AT5G47210, AT1G67430, AT3G54050, AT3G63490, AT5G02870, AT2G33450, AT3G62870, AT5G42270 and AT3G46780.

Decay modeling was performed on nuclear encoded genes with 1 TPM or more at T0 in at least one genotype. Normalized decay trajectories were modeled per gene using the RNAdecay Bioconductor package (version 1.26.0), with WT and *dcp5* as the two treatment groups and model selection by corrected Akaike information criterion (AICc). Half-life was calculated as ln(2)/α from the best-fitting model per gene. Genes were retained for downstream analysis only if the model fit passed a residual-variance quality filter (σ² < 0.0625).

### iCLIP data generation

The iCLIP experiment was performed following the protocol of Lewinski *et al*. (2024)^58^. Plants expressing DCP5 fused to YFP, or to GFP as a negative control, were grown under a 12:12 photoperiod (12 h light/12 h dark cycles, 80 μmol m⁻² s⁻¹). After two weeks, they were transferred to continuous light. Three biological replicates of DCP5-YFP-HA and two of GFP were collected per time point, at 12 and 24 hours after transfer. Samples were treated with a UV light pulse (2,000 mJ/cm²) to covalently crosslink proteins to RNA and were then immediately frozen in liquid nitrogen. Following cell lysis of the tissue, DCP5-

YFP was immunoprecipitated with GFP-Trap agarose beads (Proteintech, cat. no. gta-20). Treatment with Turbo DNase (Invitrogen) and RNase I (Invitrogen) was carried out to remove DNA in all samples and RNA in samples used for protein size reference. RNA was labeled with γ-³²P, resolved by polyacrylamide gel electrophoresis, transferred to a nitrocellulose membrane, and visualized by autoradiography. RNA of the appropriate size was recovered from the nitrocellulose membrane and used to construct libraries for sequencing. Sequencing was performed on the Illumina NextSeq 2000 platform at the Institute of Molecular Biology in Mainz, Germany.

### iCLIP data analysis

Read processing and the identification of binding peaks were carried out following the bioinformatic protocol of Lewinski *et al*. (2024)^58^. Briefly, the sequencing reads from the iCLIP experiment were sorted by biological replicate and condition, and the Illumina adapter sequences were then removed using the Flexbar package^59^. The processed reads were aligned to the TAIR10 genome with the AtRTD3 annotation using STAR^56,57^. PCR duplicates were removed using umi_tools^60^. Binding peaks were determined using PureCLIP^61^. The 2% of peaks with the worst global PureCLIP score were discarded, and from the remaining peaks the 30% with the worst score within each gene were discarded. Finally, the width of the binding sites was determined using the R package BindingSiteFinder^62^.

For the classification genes were grouped according to its biotype in protein coding (biotypes: protein coding, and non-translating CDS), long non-coding (biotypes: lncRNA and ncRNA) and short non-coding (biotypes: miRNA, snoRNA, snRNA, rRNA and tRNA). To estimate the mean distance to start and stop codons, we calculated the mean distance across isoforms for each gene. For relativize binding sites to the length of the different regions, we used the primary isoform according to Phytozome.

### Motif discovery

We searched for sequence–structure motifs enriched at DCP5 binding sites with SSMART v1.2^63^. Each binding site was extended from the center by 28 nt on either side, giving 63 nt windows; windows were trimmed at transcript ends. Identical sequences within each time point, which arise when a binding site maps to several isoforms, were collapsed into one. This left 30,254 sequences for CT12 and 71,435 for CT24.

The secondary-structure context of each sequence was predicted with RNAStructProfiling through the SSMART wrapper, which labels each nucleotide as paired or unpaired. Sequence identifiers were tagged with their time point, and the two datasets were combined into a single input of 101,689 sequences. SSMART was run with default parameters. Motifs were ranked by SSMART score, and the top-scoring motifs were reported. Position weight matrices were plotted with ggseqlogo.

### Time course sequencing experiment

For RNA sequencing, wild-type and *dcp5-1* plants were grown under a 12:12 photoperiod (12 h light/12 h dark cycles, 80 μmol m⁻² s⁻¹). After two weeks, the plants were transferred to continuous light (22 °C), and two biological replicates were harvested at 12, 16, 20, 24, 28, and 32 h in continuous light, with half of the samples treated with a 4-h pulse at 28 °C prior to harvest. Total RNA was extracted with TRIzol (Invitrogen), and libraries were prepared with the mRNA Library Prep Kit (Watchmaker). The libraries were sequenced on the Aviti platform (Element Biosciences) at the Stowers Institute for Medical Research.

### Time course sequencing experiment analysis

The RNA sequencing reads were mapped to the *Arabidopsis thaliana* TAIR10 genome^56^ using STAR 2.7.11^57^ with the following parameters: --twopassMode Basic --outFilterMultimapNmax 2 --outFilterType BySJout --outSJfilterReads Unique --sjdbOverhang 149 --alignSJoverhangMin 6 --alignSJDBoverhangMin 3 --alignIntronMin 20 --alignIntronMax 5000. Raw counts were obtained with htseq-count.

### Transcriptome rhythmicity analysis

Rhythmic genes were identified separately for each genotype with JTK_CYCLE as implemented in the meta3d function of MetaCycle (version 1.2.1)^64^, using TPM values. Genes were kept if TPM > 1 in at least 12 of the 24 22 °C libraries. Because the time series covers a single 20-h window, the six time points were concatenated to form a second cycle (CT36–CT56), and JTK_CYCLE was run on the resulting 12-time point series with the default period range (20–28 h). Genes with a Benjamini–Hochberg-adjusted q < 0.01 were called circadian clock-regulated genes (CCGs), and their phase was taken as the JTK-adjusted phase. For each genotype, relative amplitude was calculated from replicate-averaged TPM as (max − min) / (min + 1) across the six time points. CCGs with relative amplitude > 4 were called high-amplitude CCGs.

### Differential expression analysis between genotypes

Differential expression between wild-type (WT) and *dcp5* plants at 22 °C was analyzed from raw gene counts using DESeq2 (version 1.46.0)^65^. Genes were retained for downstream analyses if they had at least 10 raw counts in at least two samples, corresponding to the size of the smallest experimental group. This filtering step was performed once across all 48 libraries to ensure that the same set of genes was used in all subsequent analyses.

To assess genotype-dependent expression, two nested likelihood-ratio tests (LRTs) were performed. The genotype main effect was tested by comparing the full model ∼ time + genotype against the reduced model ∼ time. The genotype-by-time interaction was tested by comparing the full model ∼ time + genotype + time:genotype against the reduced model ∼ time + genotype. For each LRT, dispersion estimates were obtained under the corresponding full model. Genes with an adjusted P value < 0.05 in either LRT were considered differentially expressed at the global level.

To identify large, time point-specific differences, WT and *dcp5* were also compared separately at each of the six circadian times (CT12–CT32). For this, the 22 °C samples were fitted with a single factor combining genotype and time (∼ group; 12 levels), and each pairwise contrast (WT vs. *dcp5* at a given CT) was tested with a Wald test. Fitting all samples in one model let the six contrasts share size factors and dispersion estimates from the full dataset. Log₂ fold changes were shrunk with ashr (Stephens, 2017), and P-values were adjusted across genes with the Benjamini–Hochberg method within each contrast. A difference at a given time point was considered large when adjusted P < 0.05 and |log₂FC| > log₂(1.5).

Genes were subsequently classified into three categories. Robust differentially expressed genes (DEGs) were defined as genes with an adjusted P value < 0.05 in either LRT and at least one time point with an adjusted P value < 0.05 and an absolute shrunken log₂ fold change greater than log₂(1.5) in the pairwise WT-versus-*dcp5* contrast. Soft DEGs were genes significant in either LRT (adjusted P < 0.05) but without any time point meeting the per-time-point significance and fold-change criteria. Non-DEGs comprised genes that were not significant in either LRT, including genes that met the per-time-point criteria alone (n = 70).

Phase differences between genotypes were calculated only for genes that were CCGs in both genotypes, using circular arithmetic on a 24-h period: Δφ = ((φdcp5 − φWT + 12) mod 24) − 12. Antipodal values (−12 h) were set to +12 h.

### Identification of warm-temperature responsive genes

Warm-temperature responsive genes (WTRGs) were identified independently in wild type and *dcp5* using DESeq2. For each genotype, the analysis included 24 RNA-seq libraries, corresponding to six circadian time points, two temperatures, and two biological replicates per condition. Genes were classified as WTRGs if they showed a statistically significant temperature effect across the time course and an absolute temperature-dependent change greater than 1.5-fold at one or more time points.

The temperature effect was assessed using two nested likelihood ratio tests (LRTs). The first LRT compared the model ∼ time + temperature with the reduced model ∼ time, thereby identifying genes responding to temperature consistently across all time points. The second LRT compared ∼ time + temperature + time:temperature with ∼ time + temperature, thereby identifying genes for which the magnitude or direction of the temperature response depended on time of day. For each LRT, dispersion was estimated under the corresponding full model. Genes with a Benjamini–Hochberg-adjusted *P* value (*padj*) < 0.05 in either LRT were retained for subsequent time-point-level testing.

Within each genotype, a model containing 12 levels corresponding to the temperature-by-time-point combinations was fitted. The 28 °C versus 22 °C contrast was then computed at each of the six time points. Because the 28 °C treatment consisted of a 4 h pulse that ended at harvest, each contrast represents the acute transcriptional response to warm temperature during the 4 h preceding the indicated circadian time. A time point was considered significant when *padj* < 0.05 and the absolute *ashr*-shrunk log2 fold change exceeded log2(1.5). Positive log2 fold changes indicate higher expression at 28 °C than at 22 °C.

Genes satisfying both the gene-level LRT criterion and at least one significant time-point-level contrast were defined as WTRGs. This dual criterion excluded genes with statistically significant but biologically small temperature effects, as well as genes whose time-point-level response was not supported by the gene-level test. Time-point-level differential-expression calls were used in downstream analyses only for genes classified as WTRGs.

### GO term analysis

Gene Ontology (GO) enrichment of biological process terms was tested with clusterProfiler (version 4.14.6), using annotations from org.At.tair.db (version 3.20.0). Only terms annotated to at least 10 genes of the background set were tested. For each gene set, enrichment was assessed with a one-sided hypergeometric test, and p-values were adjusted for multiple testing with the Benjamini–Hochberg procedure. Terms with an adjusted p-value below 0.05 were considered significantly enriched. Fold enrichment was calculated as the proportion of genes in the set annotated to a term, divided by the corresponding proportion in the background set.

Among genes whose decay model passed σ² < 0.0625, those with a lower decay rate in *dcp5* than in wild type were classified as stabilized in *dcp5*, and those with a higher decay rate as stabilized in wild type. All 13,698 genes passing quality control served as background.

## Acknowledgements

Research was supported by the Consejo Nacional de Investigaciones Científicas y Técnicas (CONICET), the Deutscher Akademischer Austauschdienst (DAAD), the American Society of Biochemistry and Molecular Biology (ASBMB), the Agencia Nacional de Promoción de la Investigación, el Desarrollo Tecnológico y la Innovación (Agencia I+D+i), and the Deutsche Forschungsgemeinschaft (DFG, grant STA653/14-1). We acknowledge the Institute of Molecular Biology Genomics Core Facility in Mainz, Germany.

## Author contributions

Conceptualization: D.A.C., S.P.S., R.S.R., D.S., A.A.B. and M.J.Y. Data curation: D.A.C., D.J.V., V.D.A.B. Formal analysis: D.A.C., S.P.S., D.J.V., V.D.A.B. and M.L. Funding acquisition: D.A.C., S.P.S., D.S., A.A.B. and M.J.Y. Investigation: D.A.C., S.P.S., J.L.M., L.B., H.P., M.S.S., A.S.M. and M.J.d.L. Methodology: L.B., D.J.V., A.T. and M.L. Visualization: D.A.C. and S.P.S. Writing – original draft: D.A.C and M.J.Y. Writing – review & editing: D.A.C., S.P.S., J.L.M., R.S.R., D.S. and M.J.Y.

## Competing interests

The authors declare no competing interests.

## Supplementary Figures

**Supplementary Figure 1.**
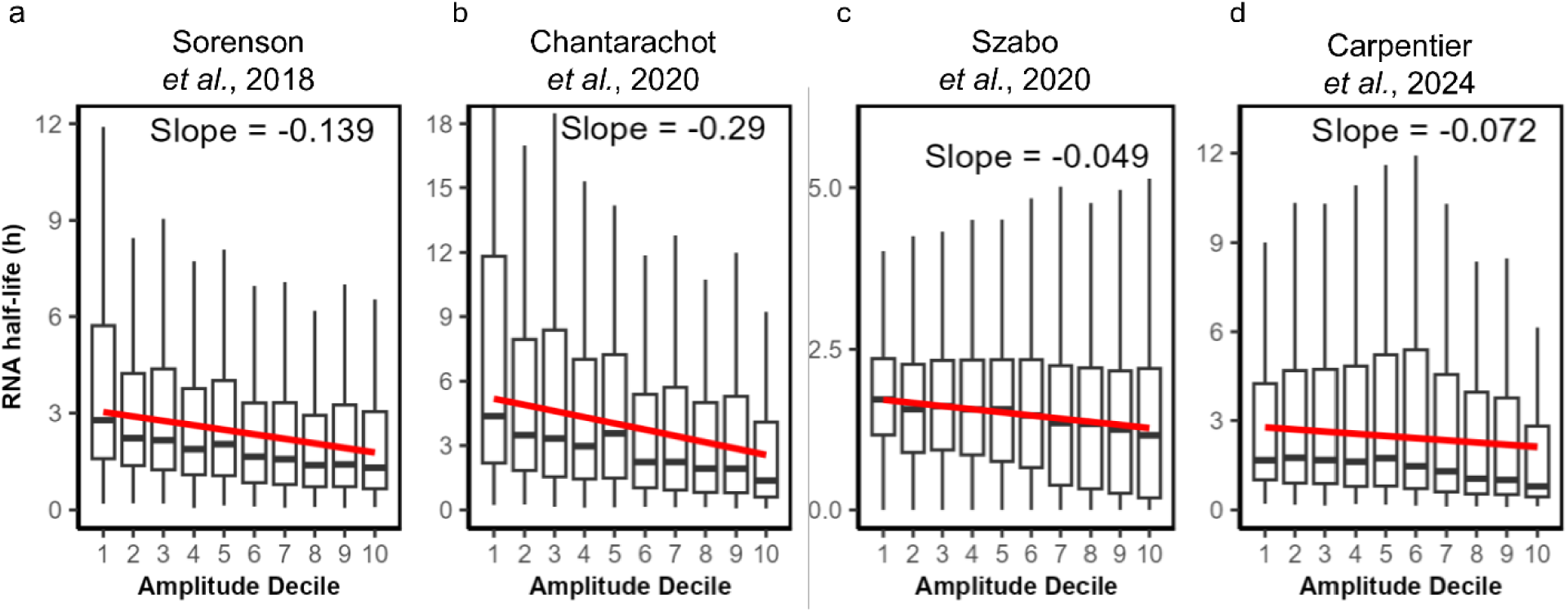
mRNA half-life decreases with circadian oscillation amplitude. CCGs were grouped into deciles of relative amplitude in each dataset (decile 1, lowest amplitude; decile 10, highest amplitude). Boxplots show the distribution of mRNA half-lives within each decile. The trendline shows a robust linear regression.

**Supplementary Figure 2.**
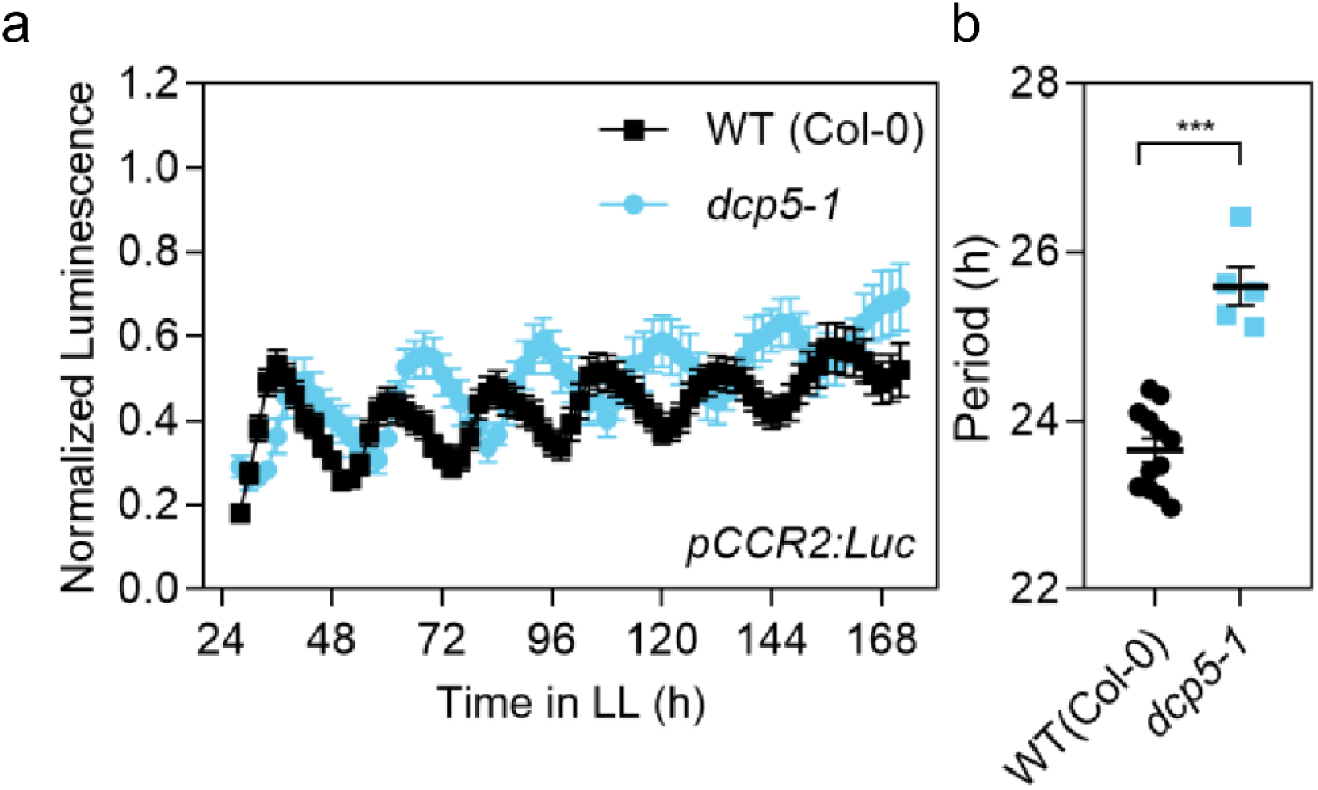
Luciferase oscillation of pCCR2:Luc. **(a)** Expression of the morning reporter pCCR2:Luc and **(b)** period length estimation. The expression was measured for 7 d in LL after entrainment under LD conditions. WT (Col-0) (n=12), *dcp5-1* mutant (n= 5). The period of bioluminescence rhythms was estimated by the fast Fourier transform nonlinear least squares (FFT-NLLS) test. Error bars represent the mean with range. ***P < 0.001 (Student’s t-test).

**Supplementary Figure 3.**
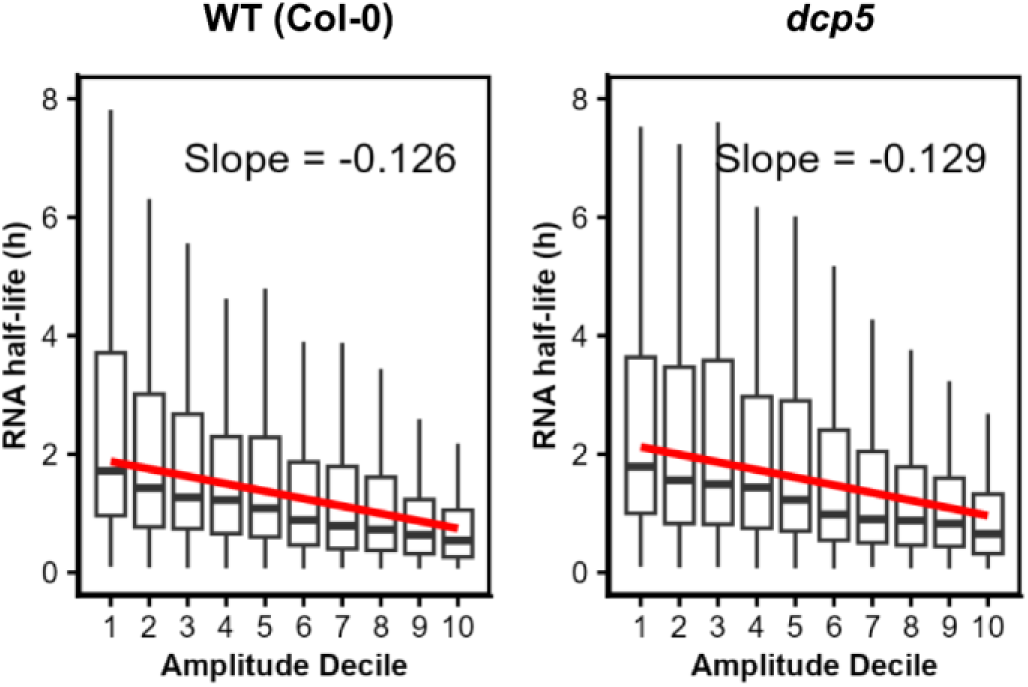
mRNA half-life decreases with circadian oscillation amplitude in both WT and *dcp5* plants. Clock controlled genes (CCG) were grouped into deciles of relative amplitude of its circadian expression (decile 1, lowest amplitude; decile 10, highest amplitude). Boxplots show the distribution of mRNA half-lives within each decile. The trendline and the slopes are the results of a robust linear regression. Relative amplitude values were taken from Romanowski *et al*. (2020).

**Supplementary Figure 4.**
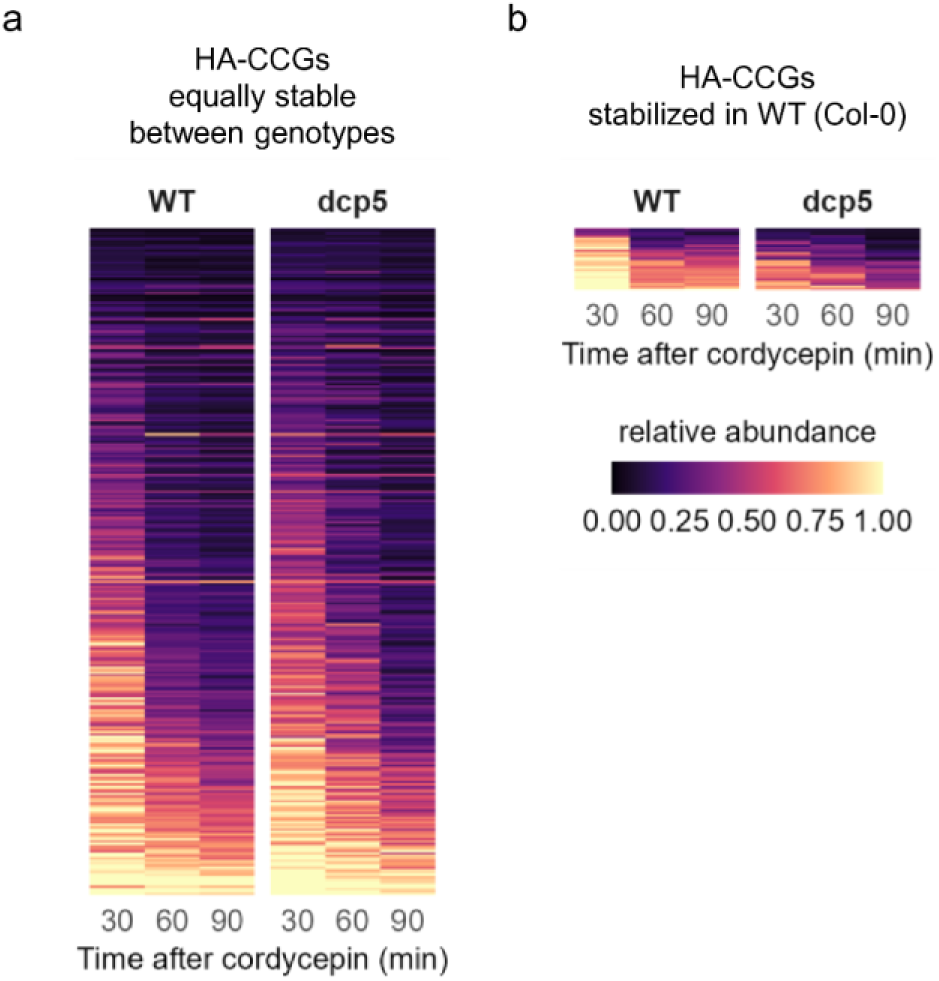
Decay profiles of high amplitude clock-controlled genes (HA-CCGs) not stabilized in *dcp5*. **a** HA-CCGs with equal decay rates in WT and *dcp5* (n = 280). **b** HA-CCGs more stable in WT than in *dcp5* (n = 26). Heatmaps show transcript abundance at 30, 60 and 90 min after cordycepin treatment, relative to t0. Rows are genes, sorted by WT half-life (shortest at the top) and kept in the same order in both genotypes.

**Supplementary Figure 5.**
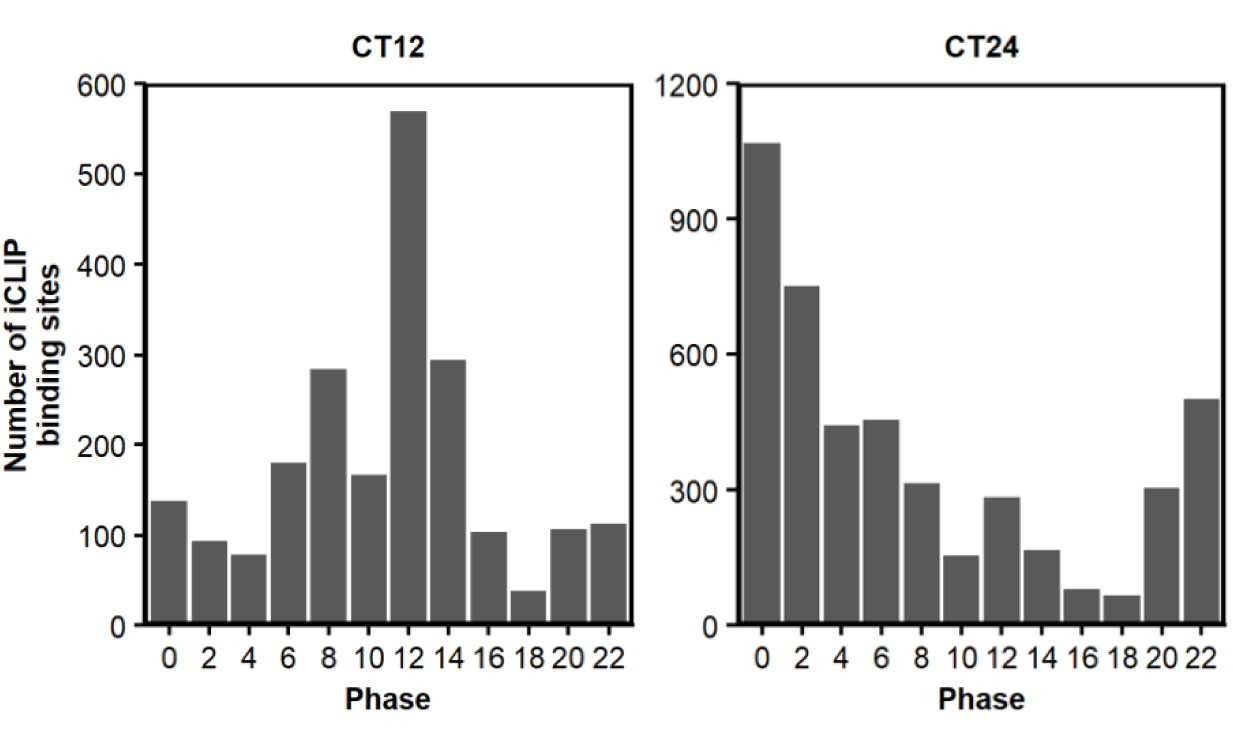
Peak-expression phases of HA-CCGs bound by DCP5. Number of DCP5 binding sites at CT12 (left) and CT24 (right) in high-amplitude clock-controlled genes (HA-CCGs; relative amplitude > 4), grouped by phase of peak expression (h). Phases and amplitudes are from Romanowski *et al*. (2020); phase 24 is plotted as 0.

**Supplementary Figure 6.**
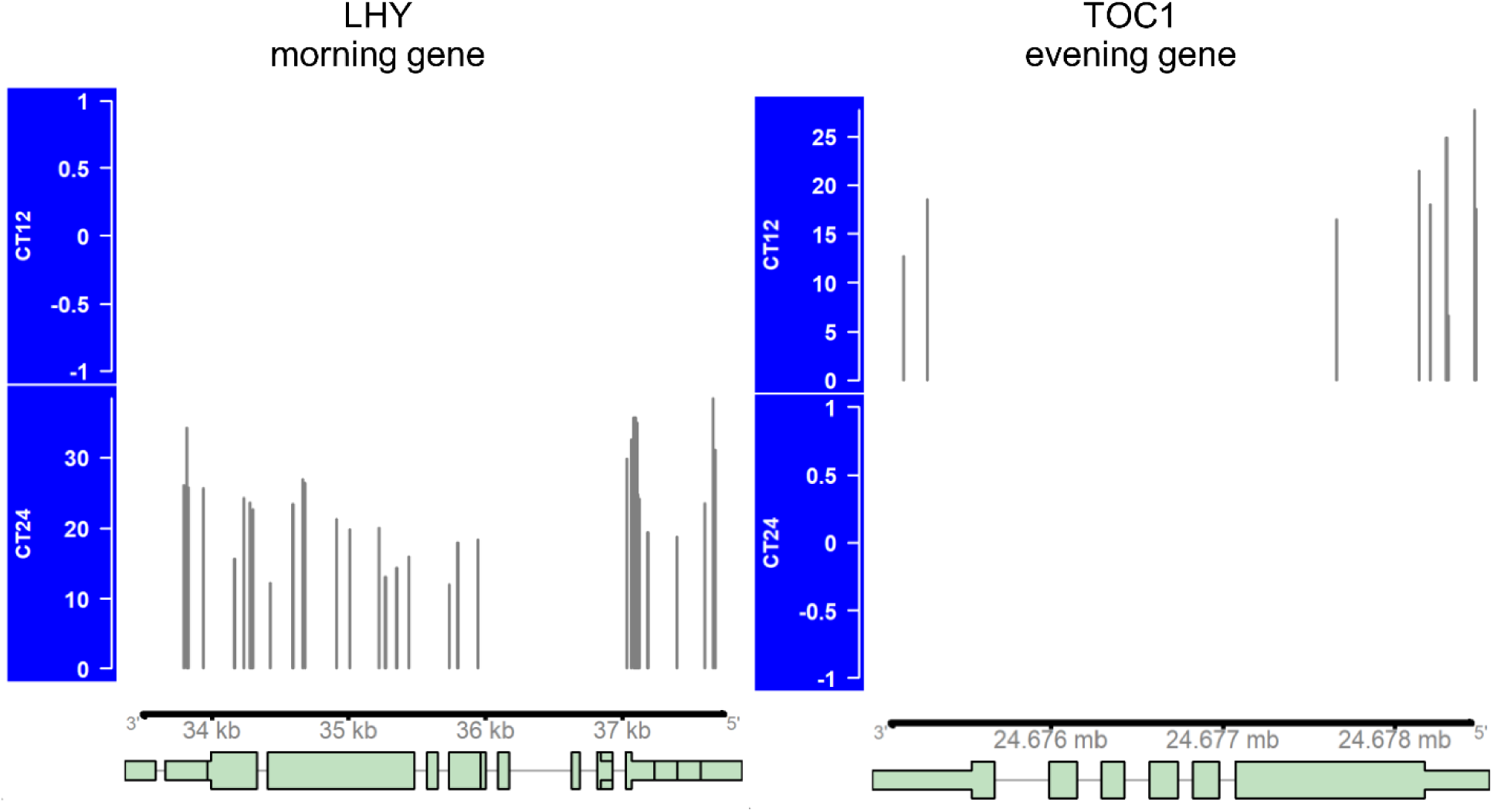
DCP5 binding sites in the dusk- and dawn-phased clock genes TOC1 and LHY. DCP5 binding sites along TOC1 (AT5G61380) and LHY (AT1G01060) at CT12 and CT24. Gene models (TAIR10) are shown below. The height of each peak represents its PureCLIP score.

**Supplementary Figure 7.**
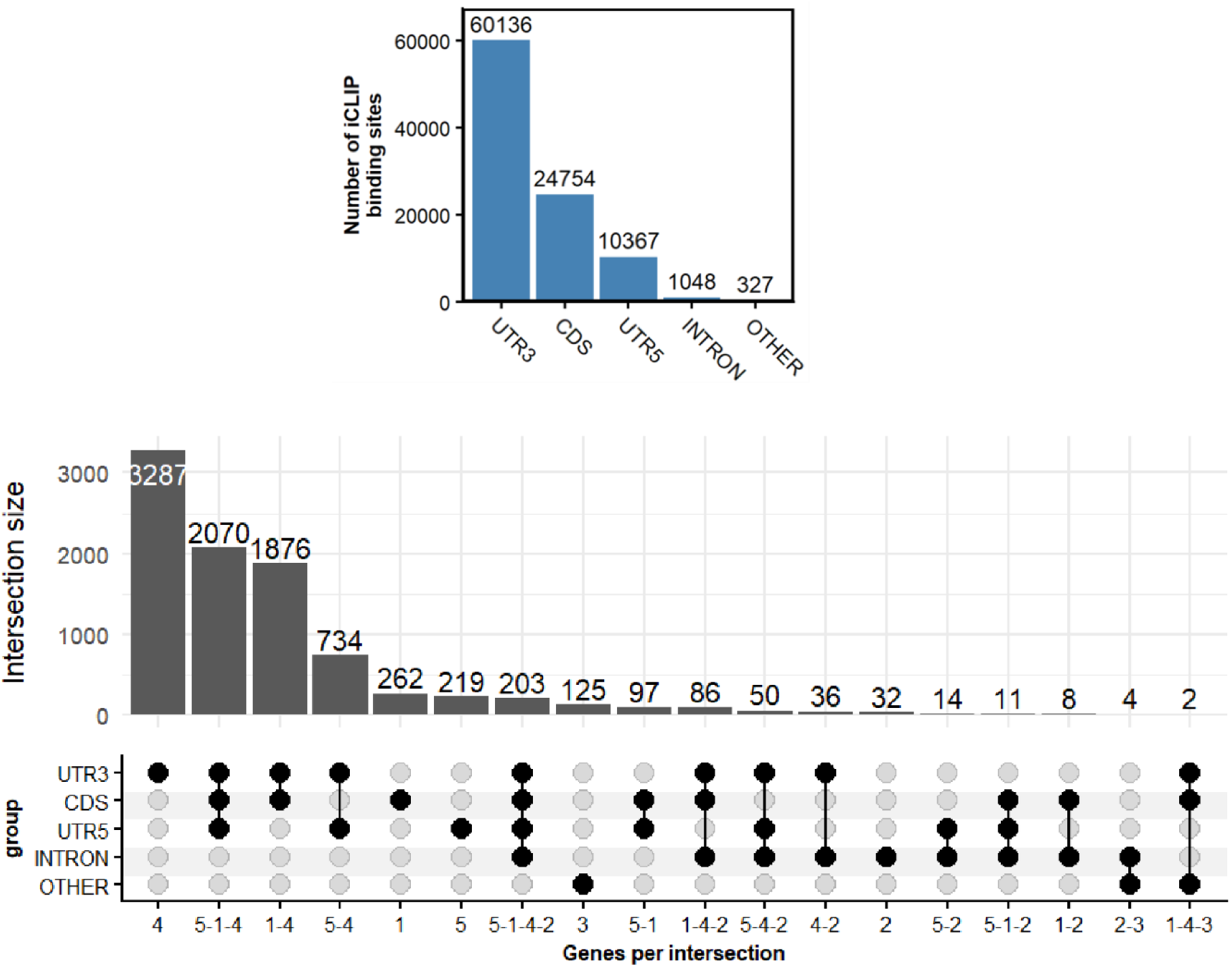
Most DCP5 binding sites are present in 3UTR. **a** Number of DCP5 binding sites in each transcript region (CT12 and CT24 combined). **b** UpSet plot of DCP5 target genes grouped by the combination of regions that contain binding sites.

**Supplementary Figure 8.**
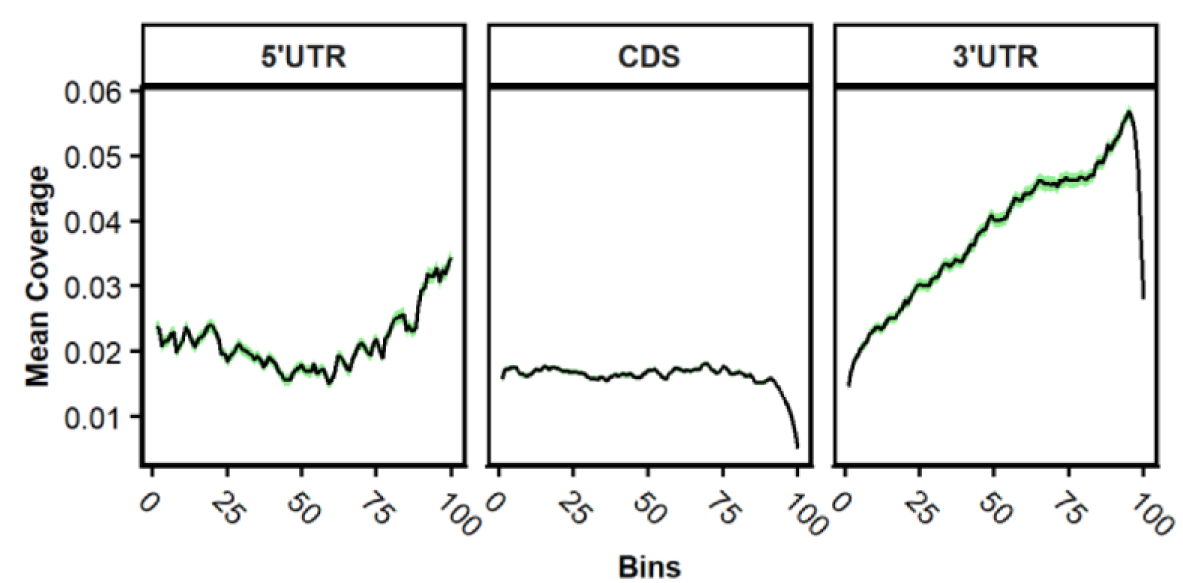
Metagene profiles of DCP5 binding across transcript regions. Mean coverage of CT24 binding sites along 5′ UTRs, CDSs and 3′ UTRs, each scaled to 100 bins.

**Supplementary Figure 9.**
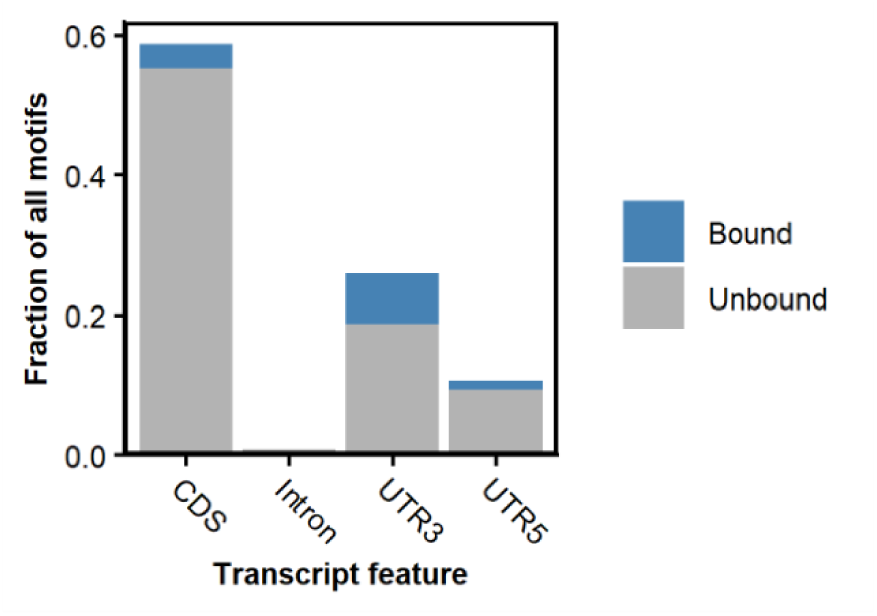
Distribution of bound and unbound motif occurrences across transcript regions. Fraction of all transcriptome occurrences of the three DCP5 motifs (WAWUW, WAMAYW and WNUAMW) found in each transcript region. Occurrences within 25 nt of a DCP5 binding site predicted by SSMART as a target of the same motif are classed as bound. (IUPAC code, W = A/U; M = A/C; Y = C/U).

**Supplementary Figure 10.**
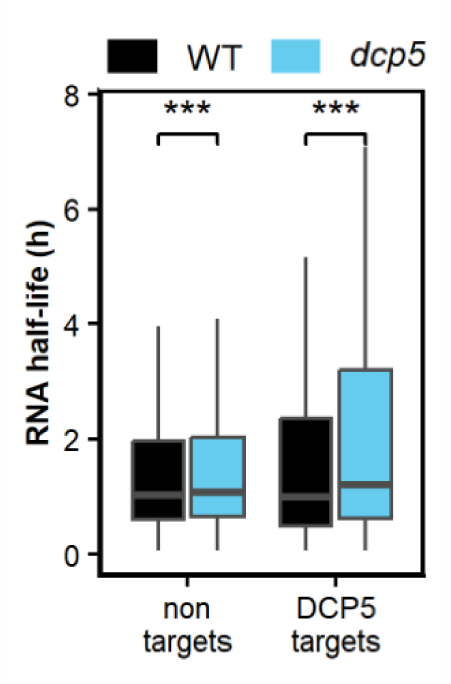
Half-lives of DCP5 target and non-target transcripts in wild-type and *dcp5* plants. mRNA half-lives of DCP5 target and non-target transcripts in WT (Col-0) and *dcp5-1*.

**Supplementary Figure 11.**
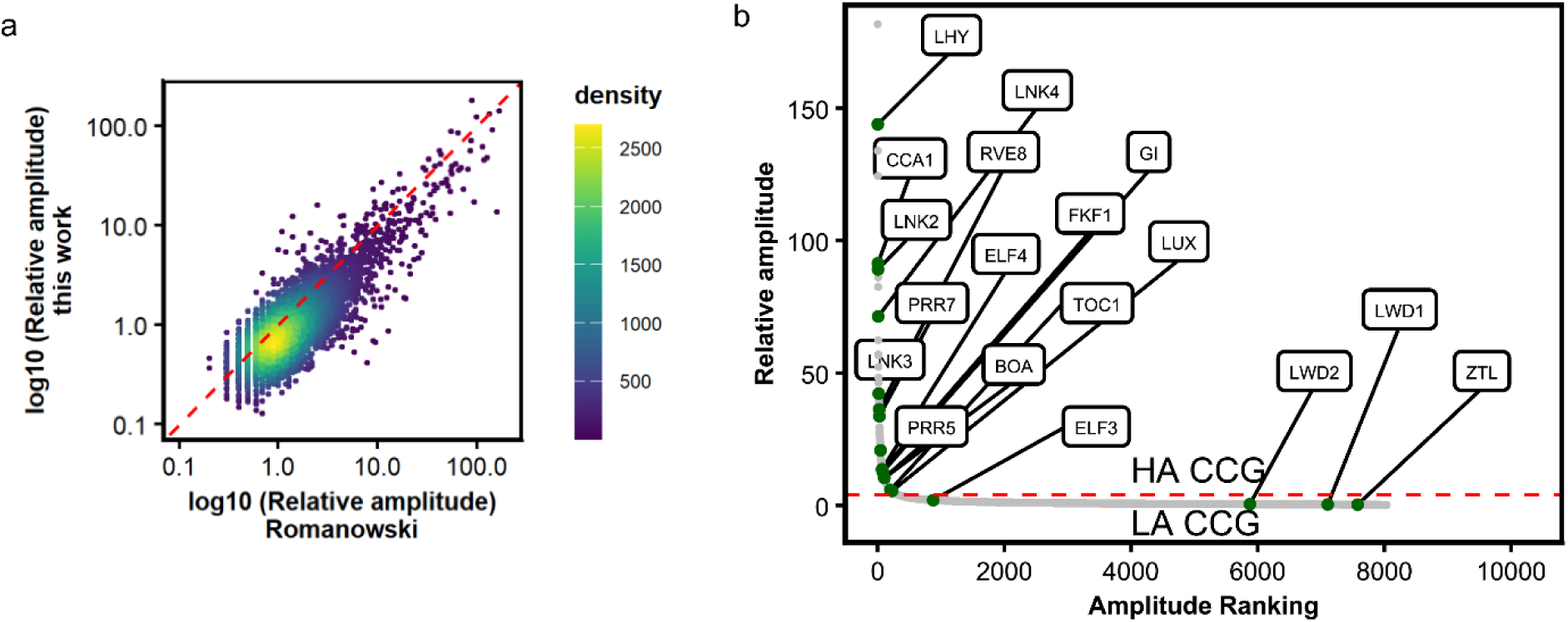
Comparison of CCG identified by Romanowski *et al.* (2020) and this study. **a** Correlation between relative amplitude in clock-controlled genes (CCG) identified by both Romanowski *et al.* (2020) and by our WT dataset. **b** Relative amplitude of the 8,036 CCGs identified in our WT dataset ranked in descending order. Core oscillator genes are labeled and highlighted in green. The red dashed line indicates the threshold at relative amplitude = 4, used to classify CCGs as high-amplitude (HA-CCGs) or low-amplitude (LA-CCGs).

**Supplementary Figure 12.**
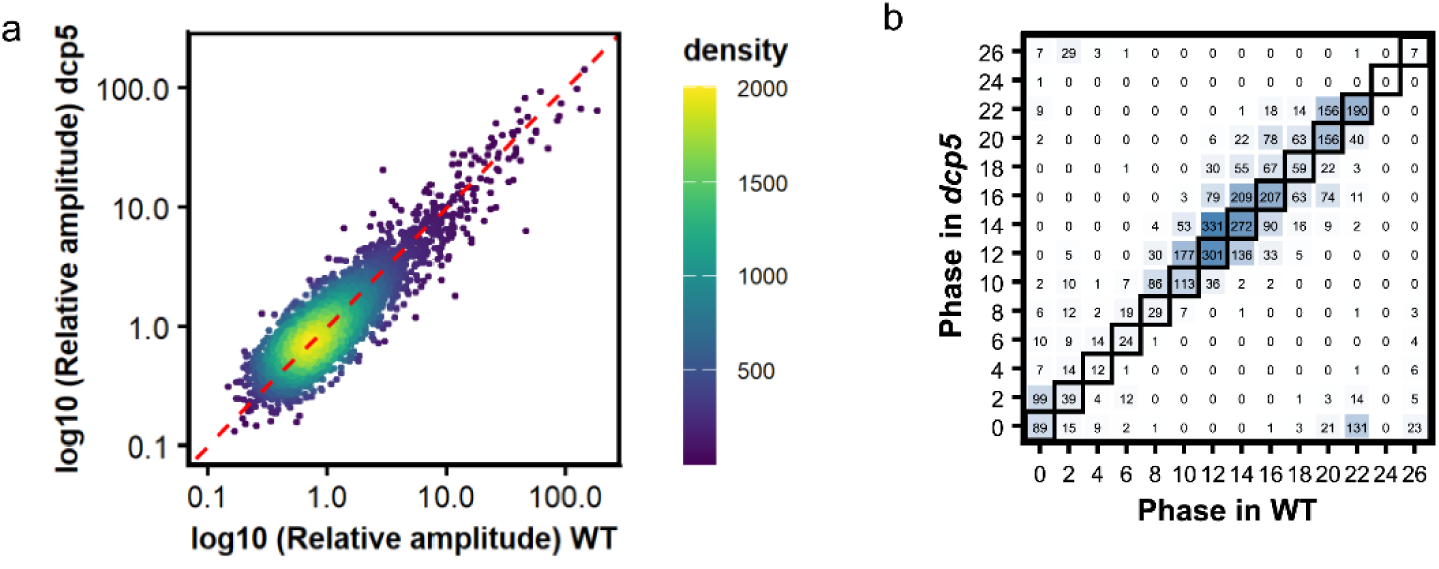
Comparison of CCG identified in wild-type and *dcp5* plants. **a** Correlation between relative amplitude in clock-controlled genes (CCG) identified in our WT and *dcp5* in our datasets. **b** Correlation plot of phase of expression in CCG found in both genotypes.

**Supplementary Figure 13.**
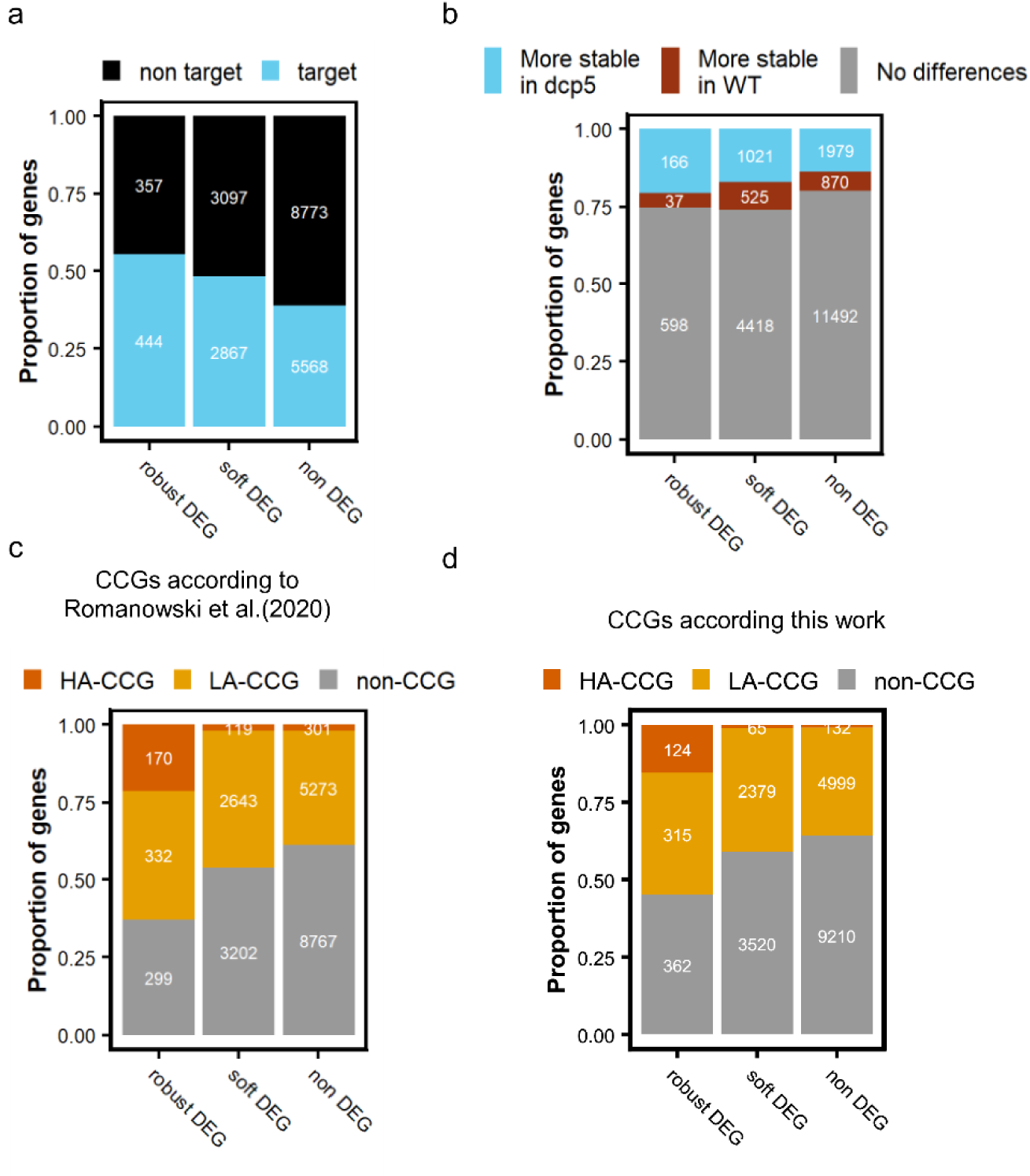
Analysis of differentially expressed genes in *dcp5* mutant plants. **a-d** Proportion of **(a)** DCP5 targets determined by iCLIP, **(b)** stabilized, destabilized, and equally stable transcripts between WT and *dcp5* determined in the cordycepin timecourse assay, high amplitude (HA-), low amplitude (LA-) and non clock-controlled genes (CCGs) in wild type plants determined by **(c)** Romanowski *et al*. (2020) and **(d)** this work among different differential expressed genes (DEG) categories. Soft DEGs have a FDR < 0.05 in the likelihood ratio test (LRT), while robust DEGs also have at least one timepoint with |log2FoldChange(*dcp5*/WT)| > log2(1.5). non-DEGs have and FDR > 0.05 in the LRT.

## Notes

### Competing Interest Statement

The authors have declared no competing interest.

## References

1. Young, M. W. & Kay, S. A. Time zones: a comparative genetics of circadian clocks. Nature Reviews Genetics 2, 702–715 (2001).

2. Nakamichi, N. The transcriptional network in the arabidopsis circadian clock system. Genes 11, 1–13 (2020).

3. Kamioka, M. et al. Direct repression of evening genes by CIRCADIAN CLOCK-ASSOCIATED1 in the Arabidopsis circadian clock. Plant Cell 28, 696–711 (2016).

4. Lu, S. X. et al. CCA1 and ELF3 interact in the control of hypocotyl length and flowering time in arabidopsis. Plant Physiology 158, 1079–1088 (2012).

5. Nagel, D. H. et al. Genome-wide identification of CCA1 targets uncovers an expanded clock network in *Arabidopsis*. Proceedings of the National Academy of Sciences 112, (2015).

6. Pérez-García, P., Ma, Y., Yanovsky, M. J. & Mas, P. Time-dependent sequestration of RVE8 by LNK proteins shapes the diurnal oscillation of anthocyanin biosynthesis. Proceedings of the National Academy of Sciences 112, 5249–5253 (2015).

7. Xie, Q. et al. LNK1 and LNK2 Are Transcriptional Coactivators in the *Arabidopsis* Circadian Oscillator. The Plant Cell 26, 2843–2857 (2014).

8. Ma, Y., Gil, S., Grasser, K. D. & Mas, P. Targeted Recruitment of the Basal Transcriptional Machinery by LNK Clock Components Controls the Circadian Rhythms of Nascent RNAs in Arabidopsis. The Plant Cell 30, 907–924 (2018).

9. Huang, W. et al. Mapping the core of the Arabidopsis circadian clock defines the network structure of the oscillator. Science 335, 75–79 (2012).

10. Nakamichi, N. et al. PSEUDO-RESPONSE ReGULATORS 9, 7, and 5 are transcriptional repressors in the Arabidopsis circadian clock. Plant Cell 22, 594–605 (2010).

11. Chow, B. Y., Helfer, A., Nusinow, D. A. & Kay, S. A. ELF3 recruitment to the PRR9 promoter requires other Evening Complex members in the Arabidopsis circadian clock. Plant Signaling & Behavior 7, 170–173 (2012).

12. Dixon, L. E. et al. Temporal repression of core circadian genes is mediated through EARLY FLOWERING 3 in Arabidopsis. Current Biology 21, 120–125 (2011).

13. Helfer, A. et al. LUX ARRHYTHMO encodes a nighttime repressor of circadian gene expression in the Arabidopsis core clock. Current Biology 21, 126–133 (2011).

14. Herrero, E. et al. EARLY FLOWERING4 recruitment of EARLY FLOWERING3 in the nucleus sustains the Arabidopsis circadian clock. Plant Cell 24, 428–443 (2012).

15. Mizuno, T. et al. Ambient temperature signal feeds into the circadian clock transcriptional circuitry through the EC night-time repressor in arabidopsis thaliana. Plant and Cell Physiology 55, 958–976 (2014).

16. Mateos, J. L., de Leone, M. J., Torchio, J., Reichel, M. & Staiger, D. Beyond Transcription: Fine-Tuning of Circadian Timekeeping by Post-Transcriptional Regulation. Genes 9, 616 (2018).

17. Zhang, X. & Guo, H. mRNA decay in plants : both quantity and quality matter. Current Opinion in Plant Biology 35, 138–144 (2017).

18. James, A. B. et al. Alternative Splicing Mediates Responses of the *Arabidopsis* Circadian Clock to Temperature Changes. The Plant Cell 24, 961–981 (2012).

19. Kwon, Y. J., Park, M. J., Kim, S. G., Baldwin, I. T. & Park, C. M. Alternative splicing and nonsense-mediated decay of circadian clock genes under environmental stress conditions in Arabidopsis. BMC Plant Biology 14, (2014).

20. Wang, X. et al. A photoregulatory mechanism of the circadian clock in Arabidopsis. Nature Plants 7, 1397–1408 (2021).

21. Perez-Santángelo, S. et al. Role for LSM genes in the regulation of circadian rhythms. Proceedings of the National Academy of Sciences 111, 15166–15171 (2014).

22. Careno, D. A., Perez Santangelo, S., Macknight, R. C. & Yanovsky, M. J. The 5ʹ-3ʹ mRNA Decay Pathway Modulates the Plant Circadian Network in Arabidopsis. Plant and Cell Physiology 63, 1709–1719 (2022).

23. Prasetyaningrum, P. et al. Inhibition of RNA degradation integrates the metabolic signals induced by osmotic stress into the Arabidopsis circadian system. Journal of Experimental Botany 74, 5805–5819 (2023).

24. Agrofoglio, Y. C. et al. Arginine methylation of SM-LIKE PROTEIN 4 antagonistically affects alternative splicing during Arabidopsis stress responses. Plant Cell 36, 2219–2237 (2024).

25. Blake, L. A., Watkins, L., Liu, Y., Inoue, T. & Wu, B. A rapid inducible RNA decay system reveals fast mRNA decay in P-bodies. Nature Communications 15, (2024).

26. Xu, J. & Chua, N.-H. *Arabidopsis* Decapping 5 Is Required for mRNA Decapping, P-Body Formation, and Translational Repression during Postembryonic Development. The Plant Cell 21, 3270–3279 (2009).

27. Schaffer, R. et al. The late elongated hypocotyl mutation of Arabidopsis disrupts circadian rhythms and the photoperiodic control of flowering. Cell 93, 1219–29 (1998).

28. Wang, Z. Y. & Tobin, E. M. Constitutive expression of the CIRCADIAN CLOCK ASSOCIATED 1 (CCA1) gene disrupts circadian rhythms and suppresses its own expression. Cell 93, 1207–17 (1998).

29. Alabadí, D. et al. Reciprocal regulation between TOC1 and LHY/CCA1 within the Arabidopsis circadian clock. Science (New York, N.Y.) 293, 880–3 (2001).

30. Gendron, J. M. et al. Arabidopsis circadian clock protein, TOC1, is a DNA-binding transcription factor. Proceedings of the National Academy of Sciences of the United States of America 109, 3167–72 (2012).

31. Romanowski, A., Schlaen, R. G., Perez-Santangelo, S., Mancini, E. & Yanovsky, M. J. Global transcriptome analysis reveals circadian control of splicing events in Arabidopsis thaliana. The Plant Journal 103, 889–902 (2020).

32. Sorenson, R. S., Deshotel, M. J., Johnson, K., Adler, F. R. & Sieburth, L. E. Arabidopsis mRNA decay landscape arises from specialized RNA decay substrates, decapping-mediated feedback, and redundancy. Proceedings of the National Academy of Sciences 115, E1485–E1494 (2018).

33. Chantarachot, T. et al. DHH1/DDX6-like RNA helicases maintain ephemeral half-lives of stress-response mRNAs. Nature Plants 6, 675–685 (2020).

34. Carpentier, M. C. et al. Genome-wide analysis of mRNA decay in Arabidopsis shoot and root reveals the importance of co-translational mRNA decay in the general mRNA turnover. Nucleic Acids Research 52, 7910–7924 (2024).

35. Szabo, E. X. et al. Metabolic Labeling of RNAs Uncovers Hidden Features and Dynamics of the Arabidopsis Transcriptome. Plant Cell 32, 871–887 (2020).

36. Gutierrez, R. A., Ewing, R. M., Cherry, J. M. & Green, P. J. Identification of unstable transcripts in Arabidopsis by cDNA microarray analysis: Rapid decay is associated with a group of touch- and specific clock-controlled genes. Proceedings of the National Academy of Sciences 99, 11513–11518 (2002).

37. Iwasaki, S., Takeda, A., Motose, H. & Watanabe, Y. Characterization of Arabidopsis decapping proteins AtDCP1 and AtDCP2, which are essential for post-embryonic development. FEBS Letters 581, 2455–2459 (2007).

38. Xu, J., Yang, J.-Y., Niu, Q.-W. & Chua, N.-H. Arabidopsis DCP2, DCP1, and VARICOSE Form a Decapping Complex Required for Postembryonic Development. The Plant Cell 18, 3386–3398 (2007).

39. Xu, J., Yang, J.-Y., Niu, Q.-W. & Chua, N.-H. Arabidopsis DCP2, DCP1, and VARICOSE form a decapping complex required for postembryonic development. The Plant cell 18, 3386–3398 (2006).

40. Narsai, R., Howell, K. A., Millar, A. H., Small, I. & Whelan, J. Genome-Wide Analysis of mRNA Decay Rates and Their Determinants in Arabidopsis thaliana. 19, 3418–3436 (2007).

41. Meyer, K. et al. Adaptation of iCLIP to plants determines the binding landscape of the clock-regulated RNA-binding protein AtGRP7. Genome Biology 18, 204 (2017).

42. Goeres, D. C. et al. Components of the Arabidopsis mRNA Decapping Complex Are Required for Early Seedling Development. the Plant Cell Online 19, 1549–1564 (2007).

43. De Alba, A. E. M. et al. In plants, decapping prevents RDR6-dependent production of small interfering RNAs from endogenous mRNAs. Nucleic Acids Research 43, 2902–2913 (2015).

44. Hoffmann, G. et al. Arabidopsis RNA processing body components LSM1 and DCP5 aid in the evasion of translational repression during Cauliflower mosaic virus infection. Plant Cell 34, 3128–3147 (2022).

45. Lidder, P., Gutiérrez, R. A., Salomé, P. A., McClung, C. R. & Green, P. J. Circadian Control of Messenger RNA Stability. Association with a Sequence-Specific Messenger RNA Decay Pathway. Plant Physiology 138, 2374–2385 (2005).

46. Batool, A., Aashaq, S. & Andrabi, K. I. Eukaryotic initiation factor 4E (eIF4E): A recap of the cap-binding protein. J of Cellular Biochemistry 120, 14201–14212 (2019).

47. Zhang, J. et al. The eukaryotic translation initiation factor eIF4E regulates flowering and circadian rhythm in Arabidopsis. Plant Journal 10.1111/tpj.16975 (2024) doi:10.1111/tpj.16975.

48. Huang, Y., McNeil, G. P. & Jackson, F. R. Translational Regulation of the DOUBLETIME/CKIδ/ε Kinase by LARK Contributes to Circadian Period Modulation. PLoS Genet 10, e1004536 (2014).

49. Chicois, C. et al. The UPF1 interactome reveals interaction networks between RNA degradation and translation repression factors in Arabidopsis. The Plant Journal 96, 119–132 (2018).

50. Jang, G.-J., Yang, J.-Y., Hsieh, H.-L. & Wu, S.-H. Processing bodies control the selective translation for optimal development of Arabidopsis young seedlings. Proceedings of the National Academy of Sciences 116, 6451–6456 (2019).

51. Xu, J. & Chua, N.-H. Dehydration stress activates Arabidopsis MPK6 to signal DCP1 phosphorylation. The EMBO Journal 31, 1975–1984 (2012).

52. Wang, Z. et al. A cytoplasmic osmosensing mechanism mediated by molecular crowding–sensitive DCP5. Science 386, eadk9067 (2024).

53. Wawer, I. et al. mRNA Decapping and 5′-3′ Decay Contribute to the Regulation of ABA Signaling in Arabidopsis thaliana. Frontiers in Plant Science 9, 1–12 (2018).

54. Choudhary, M. K., Nomura, Y., Shi, H., Nakagami, H. & Somers, D. E. Circadian profiling of the Arabidopsis proteome using 2D-DIGE. Frontiers in Plant Science 7, (2016).

55. Clough, S. J. & Bent, A. F. Floral dip: a simplified method for *Agrobacterium* - mediated transformation of *Arabidopsis thaliana*. The Plant Journal 16, 735–743 (1998).

56. Lamesch, P. et al. The Arabidopsis Information Resource (TAIR): improved gene annotation and new tools. Nucleic Acids Research 40, D1202–D1210 (2012).

57. Dobin, A. et al. STAR: ultrafast universal RNA-seq aligner. Bioinformatics 29, 15–21 (2013).

58. Lewinski, M. et al. Mapping protein–RNA binding in plants with individual-nucleotide-resolution UV cross-linking and immunoprecipitation (plant iCLIP2). Nature Protocols 19, 33–48 (2024).

59. Roehr, J. T., Dieterich, C. & Reinert, K. Flexbar 3.0 – SIMD and multicore parallelization. Bioinformatics 33, 2941–2942 (2017).

60. Smith, T., Heger, A. & Sudbery, I. UMI-tools: modeling sequencing errors in Unique Molecular Identifiers to improve quantification accuracy. Genome Res. 27, 491–499 (2017).

61. Krakau, S., Richard, H. & Marsico, A. PureCLIP: Capturing target-specific protein-RNA interaction footprints from single-nucleotide CLIP-seq data. Genome Biology 18, 1–17 (2017).

62. Busch, A., Brüggemann, M., Ebersberger, S. & Zarnack, K. iCLIP data analysis: A complete pipeline from sequencing reads to RBP binding sites. Methods 178, 49–62 (2020).

63. Munteanu, A., Mukherjee, N. & Ohler, U. SSMART: sequence-structure motif identification for RNA-binding proteins. Bioinformatics 34, 3990–3998 (2018).

64. Wu, G., Anafi, R. C., Hughes, M. E., Kornacker, K. & Hogenesch, J. B. MetaCycle: an integrated R package to evaluate periodicity in large scale data. Bioinformatics 32, 3351–3353 (2016).

65. Love, M. I., Huber, W. & Anders, S. Moderated estimation of fold change and dispersion for RNA-seq data with DESeq2. Genome Biology 15, 1–21 (2014).

